# Redox-driven control of 4-1BB interactome and signaling

**DOI:** 10.64898/2026.11.12.766337

**Authors:** Boglarka Zambo, Balint Jezso, Salla Keskitalo, Elisa Kirigin, Arnaud Landra Willm, Antti Tuhkala, Dicle Malaymar Pinar, Kari Salokas, Carole Belliardo, Bastien Morlet, Luc Negroni, Andras Zeke, Guillaume Sandoz, Markku Varjosalo, Gergo Gogl

## Abstract

4-1BB (CD137/TNFRSF9) is a major immunoregulatory receptor, yet the molecular principles governing its intracellular signaling remain unclear. Here we show that the conserved cytoplasmic cysteine residues of 4-1BB are critical determinants of its interactome, enabling redox-responsive engagement of TRAF2 and LANCL2. Using quantitative assays combined with mutational analysis, we redefine the TRAF2-binding region and show that mild oxidation induces a cysteine-dependent disulfide stabilization of the complex. We identify that 4-1BB interacts with GPX1 independently of LANCL2, most likely indirectly via TRAF2, providing local antioxidant support when glutathione (GSH) is sufficient, whereas LANCL2, whose binding competes with GSH, is favored under mildly oxidative conditions. 4-1BB signaling was found to alter cellular redox balance, in which both TRAF2 and LANCL2 sites are critical. These findings reveal that 4-1BB functions as a redox-sensitive interaction hub, switching between distinct signaling states depending on intracellular redox conditions and the cysteine-dependent remodeling of its protein interactions.

## Introduction

The co-stimulatory receptor 4-1BB (CD137/TNFRSF9) is a pivotal regulator of immune activation and a key target in cancer immunotherapy ^1^. Despite its clinical success, how its short cytoplasmic tail converts ligand engagement into intracellular signaling remains incompletely understood. 4-1BB promotes T-cell activation, proliferation, and survival ^2^, and regulates natural killer (NK) cells and myeloid populations, including macrophages, through 4-1BB-4-1BB ligand-mediated bidirectional signaling ^3–5^. Although best known for its immune functions, 4-1BB is possibly expressed at lower levels in several non-lymphoid tissues, suggesting broader physiological roles ^6,7^. From a biomedical standpoint, 4-1BB is targeted by agonistic antibodies and its cytoplasmic tail is widely used as a co-stimulatory module in engineered chimeric antigen receptors (CARs). Accordingly, the cytoplasmic signaling domain of 4-1BB is incorporated into several FDA-approved CAR-T therapies ^8,9^, including Kymriah®, Breyanzi®, Abecma®, and Carvykti® (Figure 1A), and is also used in certain experimental CAR-NK and CAR-macrophage strategies ^10,11^.

**Figure 1.**
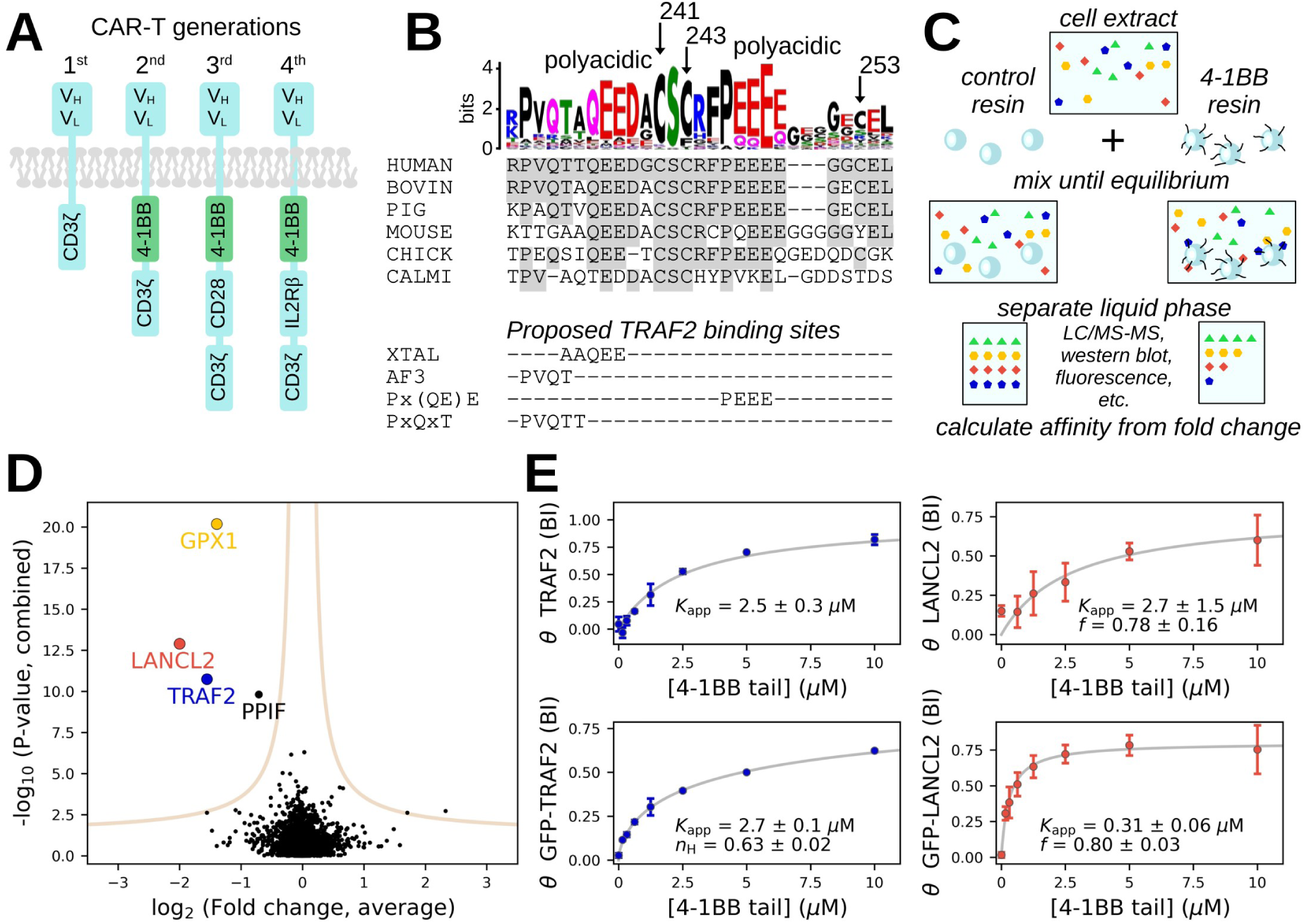
The cytoplasmic tail of 4-1BB has multiple interaction partners. (A) CAR constructs used in CAR-T therapies contain co-stimulatory modules starting from the 2^nd^ generation, such as the cytoplasmic tail of 4-1BB. Representative designs are shown from each generations. (B) Multiple residues show evolutionary conservation in the cytoplasmic tail of 4-1BB. Out of these, three distinct motifs were proposed for TRAF2 binding based on crystallography (XTAL), AlphaFold predictions (AF3), or TRAF2 consensus sequences. (C) Schematics of the native holdup (nHU) assay. (D) The cytoplasmic tailpiece of 4-1BB shows interaction in Jurkat extracts with two redox protein partners besides TRAF2: LANCL2 and GPX1. The combined results of 3 nHU experiments coupled with mass spectrometry is shown with at least 3 replicates in each measurements. (E) Titration nHU experiments quantify the apparent dissociation constants (*K*app) of endogenous or GFP-tagged TRAF2 and LANCL2, measured by Western blots from Jurkat extracts and fluorescence from transfected HEK293T cell extracts, respectively. Comparable binding was obtained between wild type and tagged TRAF2 and LANCL2 shows increased binding when tagged. Eight point titration experiments were analyzed in duplicate blots or triplicate fluorescence measurements. See supplemental Figure 1-4 for further details.

At the molecular level, 4-1BB signaling is initiated by ligand-induced receptor oligomerization ^12^, which enables recruitment of TRAF adaptor proteins, particularly TRAF2, to its cytoplasmic tail via one or more short linear motifs (SLiMs) ^13^. This interaction triggers downstream signaling cascades, including NF-κB activation ^13,14^, that eventually enhances mitochondrial biogenesis and promotes oxidative phosphorylation ^15^. This metabolic shift is accompanied by increased reactive oxygen species (ROS) production ^16,17^. Notably, CAR-T cells containing 4-1BB co-stimulatory domains similarly show enhanced mitochondrial biogenesis and oxidative metabolism compared with CD28-based CAR-T cells ^18^.

While the functional outcomes of this axis are well established, its molecular basis remains unresolved (Figure 1B). Two acidic TRAF-binding segments have been proposed within the 4-1BB tail, with the C-terminal site contributing more strongly to TRAF2 recruitment and downstream signaling ^13^. Structural studies confirmed that the N-terminal segment can associate with TRAF2 ^19^, yet calorimetric measurements reported only millimolar affinity ^20^. The same study reports motifs found in other proteins with over 30-fold stronger affinities than 4-1BB, raising the possibility that the crystallographically resolved complex captures a permissible but not necessarily physiologically dominant interaction. Indeed, crystallization can stabilize weak or transient protein-protein interactions by imposing exceptionally high local concentrations ^21^. Moreover, based on known TRAF2-binding motifs ^22^ and AlphaFold3 predictions (Supplemental Figure 1) ^23^, a third potential TRAF2 interaction site can be anticipated. Notably, the cytoplasmic tail also contains highly conserved residues outside these motifs, including multiple Cys residues that are even more conserved than the predicted TRAF2 sites, suggesting additional functional elements that remain to be elucidated.

## Results

### Proteome-wide affinity interactomic screening reveals molecular interactions mediated by the cytoplasmic tail of 4-1BB

To better define the molecular determinants of 4-1BB signaling, and characterize the interactions of the 4-1BB cytoplasmic tail, we used native holdup (nHU) approach coupled with label free quantitative mass spectrometry (MS) (Figure 1C) ^24^. Native holdup is an ideal method for capturing linear motif-mediated interactions that are often lost in conventional interactomic assays ^25^. It is a comparative chromatographic retention assay, capable not only to identify weak and highly transient molecular interactions from cell extracts, but also to simultaneously determine their apparent intrinsic binding affinities ^24,26,27^. As a bait we used a synthetic peptide corresponding the complete cytoplasmic tail of human 4-1BB (Figure 1B) and we assayed this against total Jurkat lymphoid cell extracts in three independent experiments with multiple replicates in each series. Unlike with other interactomic assays, the intrinsic properties of nHU assays allow independent experiments to be directly combined, substantially reducing the background noise of individual experiments further increasing robustness and refining resulting affinity values. As a result, we were able to confidently identify four interaction partners of the 4-1BB tail: TRAF2, LANCL2, GPX1, and PPIF (Figure 1D, Supplemental data 1, Supplemental Figure 2).

Out of these partners PPIF is a mitochondrial prolyl cis-trans isomerase, that is unlikely to interact with 4-1BB within intact cells and considered as a post lysis artifact. In contrast, TRAF2, LANCL2, and GPX1 could all potentially interact with 4-1BB in a cellular context. LANCL2, a LanC-like enzyme reported to catalyze glutathione addition to dehydroamino acids in proteins ^28^, and GPX1, a selenocysteine-containing glutathione peroxidase ^29,30^, are both involved in redox homeostasis. Based on these single point measurements, we determined dissociation constants for all three relevant partners in the range of 3-6 μM (Figure 1D) assuming simple binding mechanism. To further characterize and determine the binding mechanisms of the TRAF2 and LANCL2 interactions at higher precision, we performed titration nHU experiments and analyzed the binding of endogenous proteins with Western blot. We found that TRAF2 binds with a simple binding mechanism with an apparent dissociation constant of 2.5 μM and LANCL2 displays partial binding activity and an apparent dissociation constant of 2.7 μM (Figure 1E, Supplemental Figure 3, Supplemental Data 2). To facilitate further screening, we also measured binding of these proteins fused to green fluorescent protein (GFP) from total cell extracts prepared from transiently transfected HEK293T cells and quantified protein concentration based on fluorescence. We found that GFP-TRAF2 binding displays slight negative cooperation with an apparent dissociation constant of 2.7 μM and that GFP-LANCL2 still displays comparable partial activity as the endogenous protein, but displays a significantly stronger binding with a dissociation constant of 0.3 (Figure 1E) μM. Remarkably, these experiments found affinities 400-fold stronger between 4-1BB and TRAF2 compared to previously measured ∼1 mM affinity, found by calorimetric measurements between the motif binding domain of TRAF2 and a peptide motif of mouse 4-1BB ^20^, which places 4-1BB among the strongest reported interaction partners of TRAF2 (Figure 1E).

GPX1 has previously been reported as a TRAF2-interacting protein ^31^, and our data further clarify the nature of this interaction in the context of 4-1BB. Based on the measured fraction-bound values in nHU assays together with absolute proteomic abundance estimates, we observed an apparent binding stoichiometry in which for every 4-1BB bound TRAF2 molecule approximately 7 molecules of GPX1 is also engaged (Supplemental data 1). This ratio is somewhat consistent with a model in which the 4-1BB cytoplasmic tail is engaged with TRAF2 that is tightly bound to the tetrameric GPX1 in either 1:4 or in 1:8 stoichiometric ratios and the small discrepancy between the expected and the measured 1:7 ratio is likely due to the inaccuracy of absolute quantification by proteomics. Accordingly, our data support an indirect association of GPX1 with 4-1BB. Further experimental validation of this interaction was limited by the lack of suitable GPX1 antibodies and by the inability to express GPX1 as a GFP fusion in mammalian cells due to its central selenocysteine (Supplemental Figure 4). Mutation of the selenocysteine residue to cysteine in the GFP fusion construct did not yield detectable interaction with 4-1BB (Supplemental Figure 4), although this mutation is known to produce an enzyme with significantly reduced catalytic activity ^32^.

### Binding site mapping on 4-1BB

We performed nHU–MS assays using a series of 4-1BB mutants to map interaction determinants. These included single cysteine to serine substitutions (C241S, C243S, C253S), a mutant converting the conserved CSC motif to STS, abolishing both 241 and 243 cysteines, and a clinically observed somatic mutation, P246S, identified in skin cancer patients (Figure 2A, Supplemental Figure 5, Supplemental data 1) ^33^. Analysis of these variants revealed distinct binding requirements for different interaction partners. Strong TRAF2 binding was observed in all variants retaining cysteine 243, while C243S showed markedly weaker binding and STS a complete loss of TRAF2 binding. LANCL2 binding was selectively abolished by the mutation of cysteine 253 (Figure 2A). GPX1 binding was found to be highly variable between experimental series, suggesting a possible strong condition dependency. However, in one dataset where GPX1 showed robust binding to the wild type (WT) 4-1BB, the interaction was lost by the STS mutation, which otherwise preserved only LANCL2 binding, suggesting that these redox partners bind independently.

**Figure 2.**
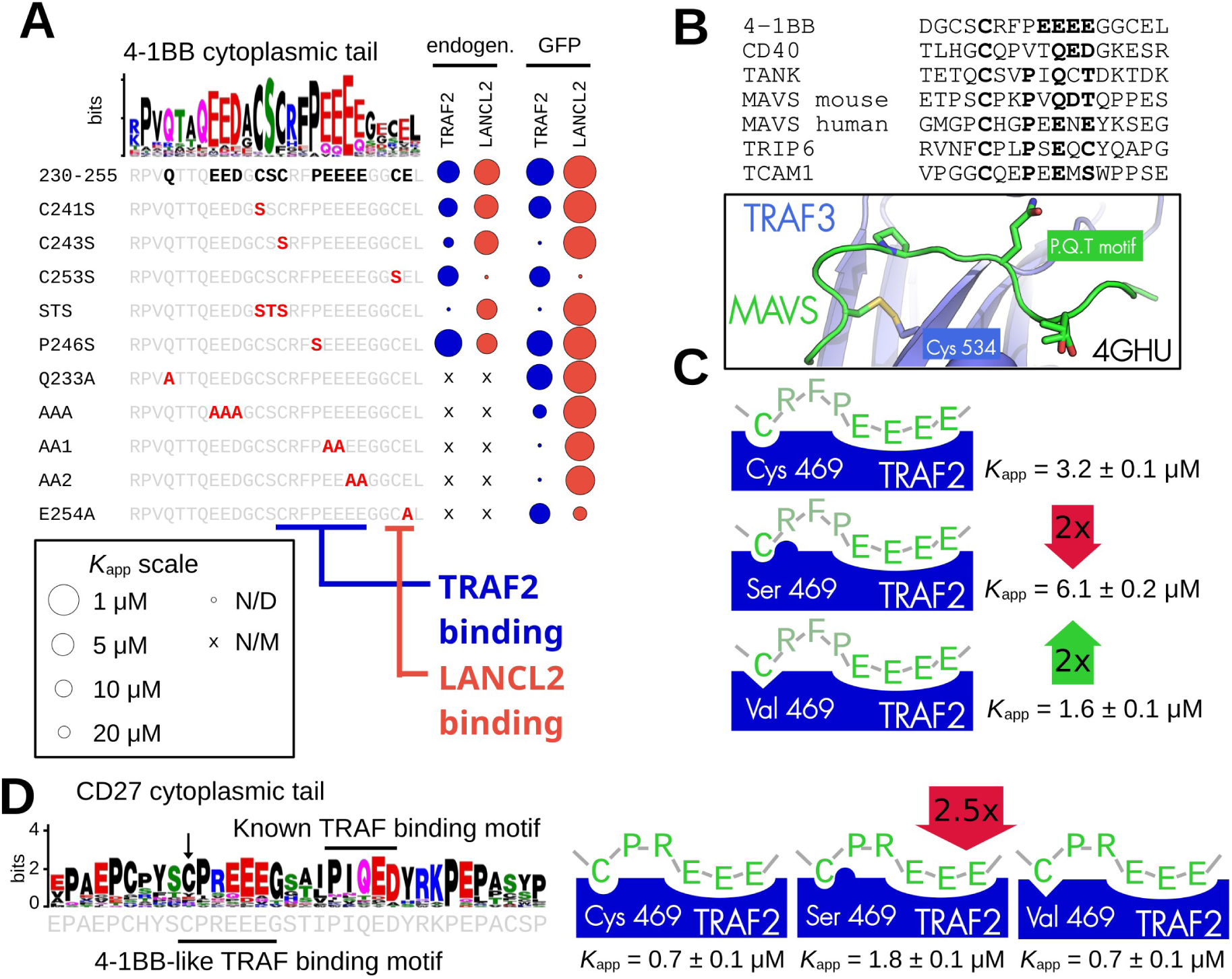
Locating binding sites on 4-1BB. (A) An array of mutant 4-1BB cytoplasmic tail peptides were used in nHU experiments to identify binding sites for partners. This revealed that TRAF2 binds mostly to the C-terminal acidic region with C243 showing a critical contribution and that LANCL2 binds to the extreme C-terminus with C253 playing a key role. Dot size is proportional to apparent dissociation constants. N/M: interaction not measured. N/D: interaction is not detected. (B) Multiple other TRAF interaction partners contain a cysteine residue preceding the main TRAF binding site. In a previously determined crystal structure, this residue was found to form a disulfide bond with a solvent exposed cysteine on TRAF3 while in a second structure it was found to only provide hydrophobic contact. (C) Mutation of the equivalent Cys residue on TRAF2 to hydrophilic serine decrease 4-1BB binding, while mutating it to hydrophobic valine increases binding, indicating a hydrophobic pocket. (D) The cytoplasmic tail of CD27 contains a conventional and a 4-1BB-like TRAF binding motif with an upstream Cys residue. The hydrophilic Ser 469 surface mutant TRAF2 shows weaker binding to CD27, similarly to 4-1BB. See supplemental Figure 5-7 for further details.

TRAF5, a homolog of TRAF2, was detected as a 4-1BB interactor in one experimental batch (batch 11), with the variants C243S, C253S, P246S, albeit with a much weaker apparent affinity compared to TRAF2 (Supplemental Figure 5C). However, no binding could be detected with extracts containing GFP-TRAF5 indicating that this interaction is indirect (Supplemental Figure 6A, Supplemental Data 2). To test this, we co-expressed GFP-TRAF5 with CFP-TRAF2 and performed nHU experiments on extracts derived from these cells. In this context, both TRAF2 and TRAF5 were detected as 4-1BB interactors (Supplemental Figure 6C, Supplemental Data 2). Such co-dependent binding was not observed when TRAF5 was co-expressed with CFP-LANCL2. This indicates that TRAF5 is recruited to 4-1BB indirectly through hetero-oligomerization with TRAF2 rather than binding directly to 4-1BB.

Several mutant 4-1BB peptides also showed weak interactions with the PDZ domain-containing proteins NHERF1 and PARD3 (Supplemental Figure 5C). Among these, PARD3 binding was also detected in one nHU–MS experiment performed with the WT 4-1BB peptide. However, no direct binding between 4-1BB and GFP-PARD3 could be observed in nHU assays using cell extracts from GFP-PARD3 expressing cells (Supplemental Figure 6B, Supplemental Data 2). We therefore assumed that, similarly to TRAF5, PARD3 might be recruited indirectly through TRAF2 or LANCL2, yet only extremely weak binding was detected in the presence of either protein (Supplemental Figure 6C, Supplemental Data 2). We also note that DCAF12 was binding to all four baits of the batch 11 experimental series that is most likely an experimental artifact as this C-terminal degron receptor may bind to the free peptide C-terminus. Additional partners appeared in individual interactomes, such as TXNDC5, that showed binding to the C241S variant, further indicating the link between 4-1BB and redox processes.

To gain further insight into the binding determinants of key 4-1BB interactions, we performed titration nHU experiments with fluorescent readout using an expanded panel of mutant 4-1BB peptides and extracts containing GFP-fused TRAF2 and LANCL2 (Figure 2A, Supplemental Figure 5A-B, Supplemental Data 2). To target the two polyacidic regions, we replaced the N-terminal EED sequence to AAA and we replaced the C-terminal EEEE sequence to AAEE (AA1) or EEAA (AA2). Mutations introduced into either of the two polyacidic regions significantly weakened TRAF2 binding compared to WT 4-1BB, with the C-terminal acidic patch exerting a stronger effect where either AA1 or AA2 completely abolished binding, consistently with an early study that first described this interaction nearly three decades ago ^13^. In contrast, a Q233A mutation engineered to disrupt the putative TRAF2-binding motif matching the PxQxT consensus, that is also supported by AlphaFold models, as well as the P246S mutation that is supposed to disrupt the motif following a Px(QE)E consensus had no detectable effect on TRAF2 binding and even the AAA mutation caused only ∼6-fold affinity decrease that seemed to be critical in the crystal structure of the mouse complex. Consistent with the nHU-MS results, C243S markedly reduced TRAF2 binding, whereas the other cysteine substitutions (C241S, C253S) had either no effect or at most a 2-fold change in affinity (Figure 2A). Surprisingly, the effect of C243S was comparable to that of the C-terminal polyacidic patch mutation. Collectively, these results demonstrate that TRAF2 engages 4-1BB through a binding mode that differs from previous reports, with both the C243 residue and the C-terminal acidic patch playing critical roles in the interaction.

For LANCL2, titration experiments clearly outlined a binding site at the extreme C terminus of 4-1BB. The interaction was completely abolished by the C253S mutation, while the adjacent E254A substitution also reduced binding affinity by 50-fold. Mutations within the upstream acidic patch clearly weakened the interaction by ∼6-fold,confirming that LANCL2 primarily engages the terminal GCE motif of 4-1BB (Figure 2A), which lies immediately downstream of the TRAF2-binding site.

### A new binding pocket on TRAF2 coordinates 4-1BB binding

The importance of Cys 243 on 4-1BB raises the question about how it contributes to TRAF2 binding. Since all tested structure prediction tools, including AlphaFold, failed to capture the contribution of this residue to the interaction, we examined other known TRAF interactions. We found that similarly positioned cysteine residues adjacent to canonical TRAF-binding motifs are also present in TANK, CD40, and mouse MAVS (Figure 2B) ^22^. In TANK, a crystal structure captured this cysteine facing a solvent-exposed cysteine of TRAF3 ^34^, Cys 534, which is analogous to Cys 469 in TRAF2 (Supplemental Figure 7), while in crystal structure of the MAVS complex the equivalent cysteine in the peptide was found oxidized with the adjacent cysteine on TRAF ^35^ (Figure 2B). These observations suggest that cysteines positioned opposite TRAF-binding motifs can engage either in hydrophobic contacts under reducing conditions or in intermolecular disulfide bond formation under oxidative conditions.

To test whether Cys 469 of TRAF2 contributes to 4-1BB binding through hydrophobic interactions, we introduced C469S and C469V mutations into full-length TRAF2 and quantified 4-1BB binding using nHU. Substitution with serine, which introduces a polar hydroxyl group, reduced binding by approximately two-fold, whereas replacement with the slightly bulkier and more hydrophobic valine increased binding to a similar extent (Figure 2C, Supplemental Figure 7, Supplemental Data 2). These results demonstrate that Cys 469 does contribute to 4-1BB recognition predominantly through hydrophobic contacts under a reducing environment, and indicate that solvent-exposed cysteines on both TRAF2 and 4-1BB strengthen the interaction. This direct interaction also signifies that under oxidative condition, this noncovalent interaction could turn covalent, similarly to the observed MAVS complex. Although the disulfide itself was not directly detected, the requirement for Cys243 (4-1BB) and Cys469 (TRAF2) and the analogy with the oxidized MAVS motif complex strongly support this model.

To further investigate this oxidation-sensitive Cys 469-mediated exosite on TRAF2, we investigated the binding of the cytoplasmic tail of CD27, another TNF receptor family member known to be capable of binding TRAF proteins ^36^. CD27 contains two putative TRAF-binding motifs, one matching the PxQE consensus sequence and a second resembling the 4-1BB motif with a polyacidic segment and a downstream Cys residue ^37^. We measured the binding of GFP-TRAF2 and its C469S and C469V variants to this synthetic peptide, comprising both motifs (Figure 2D, Supplemental Data 2). This experiment revealed that CD27 binds much tighter to TRAF2 than 4-1BB and that the C469V mutation does not affect this binding. However, by introducing the hydrophilic residue at that site via the C469S mutation, a 2.5-fold weaker binding was observed (Figure 2D). This shows that this exosite is utilized not only by 4-1BB, but also by several other TRAF interaction partners.

### Heterologous signalosome reconstruction assay to validate 4-1BB interactions in cells

Overexpression of TRAF proteins has previously been shown to induce punctate signaling assemblies ^38–40^. Similarly, overexpression of 4-1BB in HEK293T cells has previously been shown to induce similar TRAF-containing structures consistent with signalosome formation ^41,42^. To investigate cellular complex formation between 4-1BB, TRAF2, and LANCL2, we performed co-localization experiments in HEK293T cells transiently expressing fluorescently tagged constructs (4-1BB–mCherry, GFP/CFP-TRAF2, GFP/CFP-LANCL2) (Figure 3). Expression of TRAF2 in this heterologous cell line induced the formation of grape-shaped, cytoplasmic structures consistent with previously described signalosome-like assemblies ^39,41,42^. 4-1BB was greatly enriched on the periphery of these signalosomes, but it also showed co-localization with TRAF2 at discrete plasma membrane patches (Figure 3A). In contrast, LANCL2 displayed predominantly nuclear localization and showed no co-localization with 4-1BB, consistent with previous reports indicating that its association with the plasma membrane depends on N-terminal myristoylation (Figure 3B) ^43^. Upon co-expression of LANCL2 with TRAF2, LANCL2 partially relocalized to the TRAF2 signalosomes, regardless of the presence of 4-1BB (Figure 3C). These signalosomes do not show co-localization with common endosomal markers, such as Rab11, or Rab5A that was previously shown before to interact with TRAF2 (Supplemental Figure 8) ^44^. When all three partners are present, robust co-localization could be observed at the periphery of TRAF2 signalosomes and at the plasma membrane patches where TRAF2 and occasionally LANCL2 were also recruited (Figure 3D). This co-localization pattern was consistent regardless of whether CFP-TRAF2 and GFP-LANCL2 or GFP-TRAF2 and CFP-LANCL2 were used in the experiments, excluding spectral overlap artifacts.

**Figure 3.**
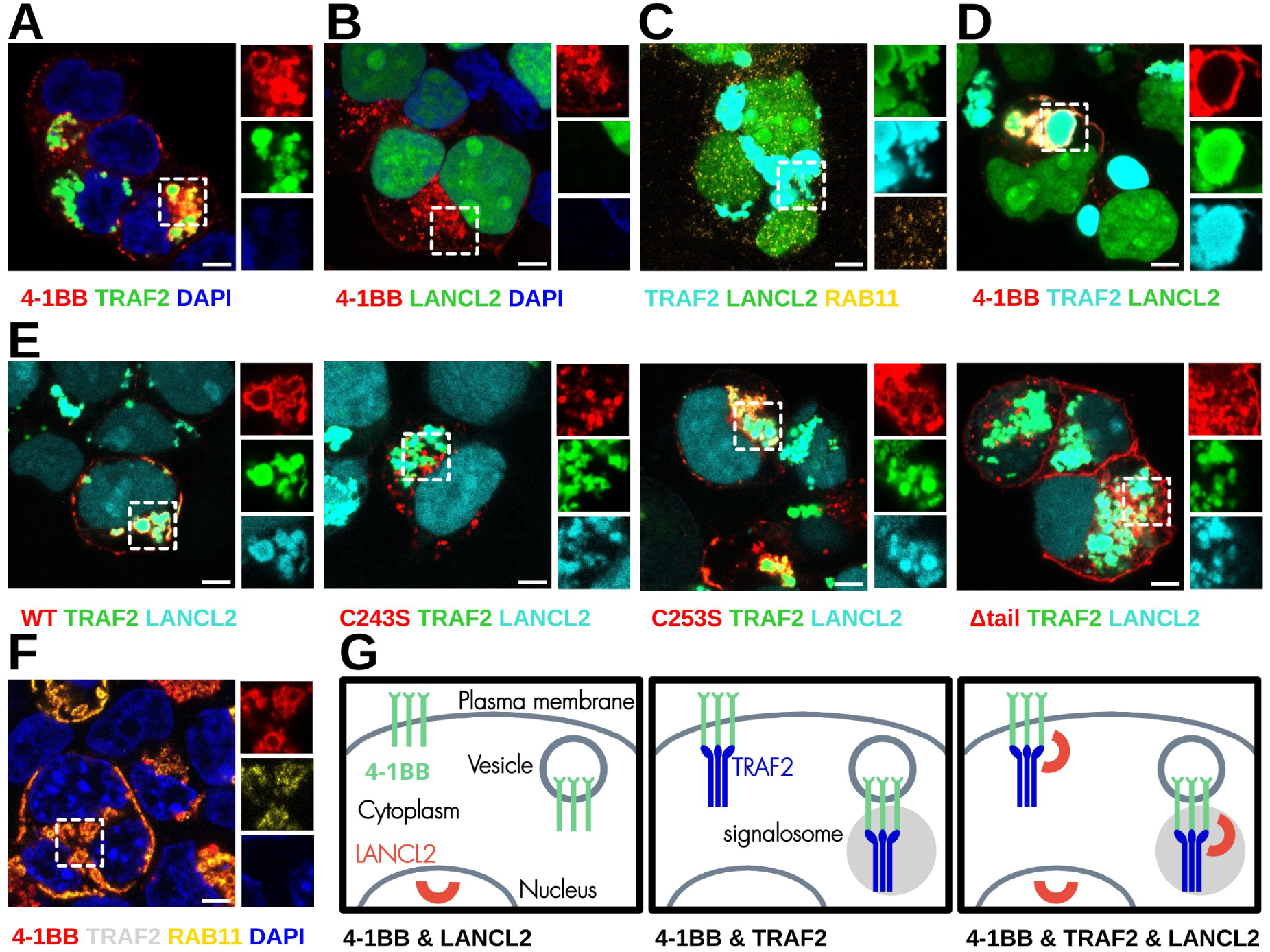
4-1BB, TRAF2, and LANCL2 show similar cellular localization. Confocal images were taken from fixed HEK293T cells transfected with fluorescently tagged proteins. (A-B) Fluorescence images of 4-1BB and TRAF2 or LANCL2. DAPI marks nucleus. (C) Fluorescence images of TRAF2 and LANCL2. RAB11 marks recycling endosomes. (D-E) Fluorescence images of different 4-1BB variants with TRAF2 or LANCL2. (F) Fluorescence image of 4-1BB in the presence of untagged TRAF2. RAB11 marks recycling endosomes and DAPI marks nucleus. (G) Schematic summary of observations. Red, green, or cyan colors indicate mCherry, eGFP, or CFP tags, respectively. DAPI staining is shown in blue. Endogenous RAB5A staining is shown in yellow. For each panel, an overlay signal is shown on the left and the insets on the right show separate channels. Scale bar is equivalent to 5 μm. Representative images are shown based on hundreds of observations. See supplemental Figure 8 for further details.

Mutation analysis revealed a critical role for C243 in this spatial organization. The C243S mutant showed a strong reduction in plasma membrane clustering and, notably, cytoplasmic 4-1BB vesicles were no longer in the proximity of TRAF2 signalosomes, indicating impaired TRAF2 association (Figure 3E). In contrast, the C253S mutant was indistinguishable from wild-type 4-1BB in both membrane localization and association with TRAF2 signalosomes (Figure 3E). As a negative control, a cytoplasmic tail–deleted 4-1BB construct displayed uniform plasma membrane distribution without clustering and localized to cytoplasmic vesicles. This 4-1BB-Δtail construct failed to recruit TRAF2 to the plasma membrane, and its vesicular structures showed no co-localization with TRAF2-signalosomes (Figure 3E). Even though TRAF2 signalosomes were negative for the tested endosomal markers, we assumed that internalized 4-1BB must be present in vesicles. Indeed, 4-1BB showed clear co-localization with Rab11, indicating that internalized 4-1BB is actively recycled in HEK293T cells (Figure 3F, Supplemental Figure 8).

### Binding mechanism of the interaction between 4-1BB and LANCL2

To further investigate the physical interaction between LANCL2 and 4-1BB and show that it is direct and stoichiometric, we performed single molecule pulldown (SiMPull) assay ^45–48^. We found strong enrichment of GFP-LANCL2 on slides coated with biotinylated WT or C243S mutant 4-1BB compared to the C253S mutant (Figure 4A, Supplemental Data 3). To estimate the binding stoichiometry of the interaction, we counted the number of discrete photobleaching steps of individual bound GFP-LANCL2 molecules. The vast majority of the observed spots showed single step fluorescence decay in photobleaching experiments for both WT and C243S mutant 4-1BB, indicating a 1:1 binding stoichiometry (Figure 4B, Supplemental Data 3).

**Figure 4.**
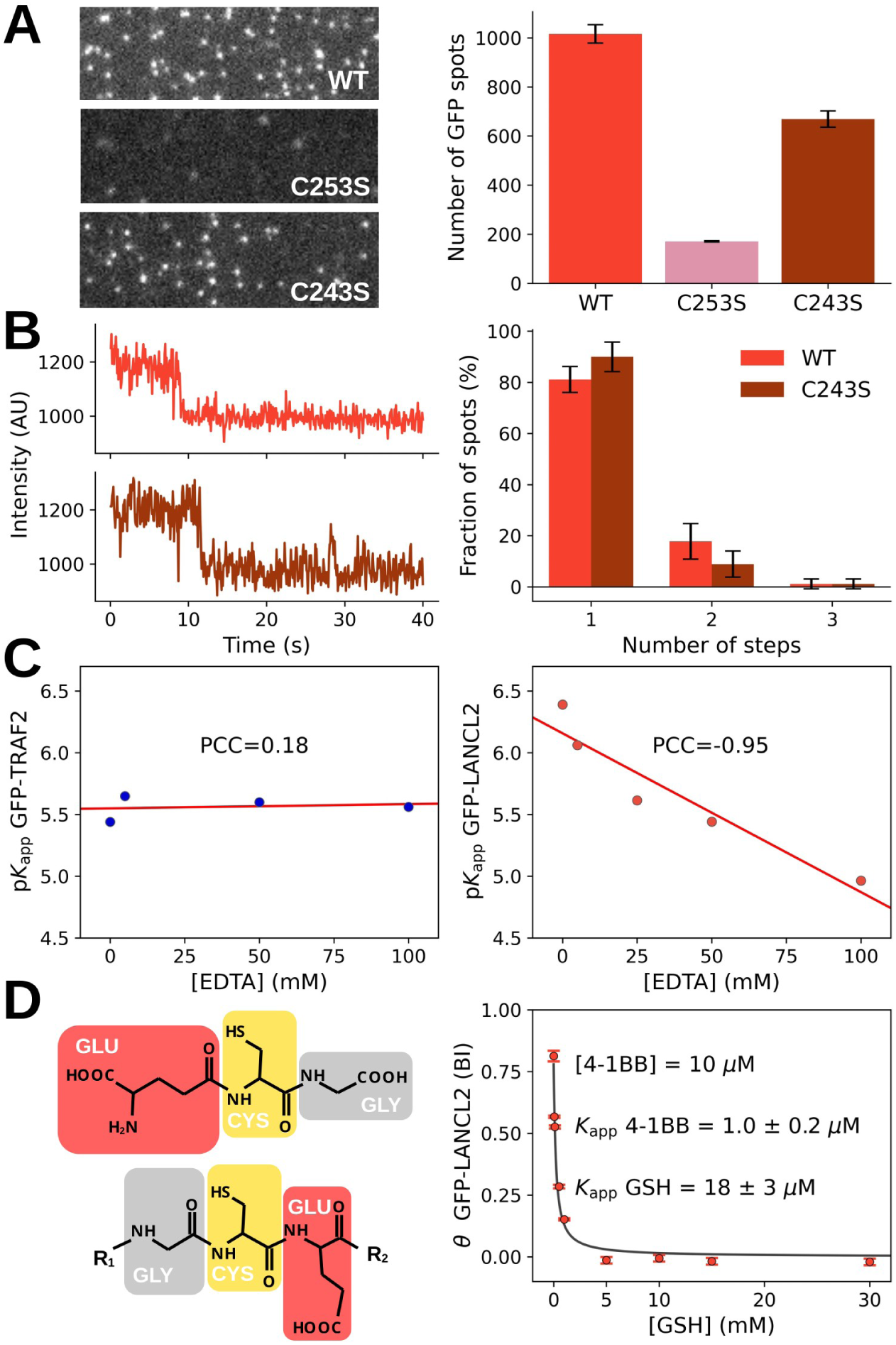
4-1BB binds to the active site of LANCL2. (A) SimPull experiments carried out with GFP-LANCL2 and surface-immobilized 4-1BB tail peptide variants. While WT and C243S mutant 4-1BB tails recruited large number of GFP-LANCL2 molecules on the surface, while the C253S variant only resulted in small amount of fluorescent spots, i.e. background signal. Representative images are shown. (B) Photobleaching traces of a GFP-LANCL2 molecule bound to a surface immobilized WT or C243S 4-1BB tail confirms 1:1 binding between 4-1BB and LANCL2. As shown on the representative traces on the left, the vast majority of GFP-LANCL2 spots showed a single-step photobleaching. (C) The removal of the bound Zn^2+^ ion from the RING domain of TRAF2 by supplementing nHU reactions with EDTA does not affect its interaction with 4-1BB (left), while the removal of the Zn^2+^ ion from the active site of LANCL2 by EDTA strongly impairs 4-1BB binding (right). (D) The natural ligand of LANCL2, GSH resembles the reversed sequence of the LANCL2 binding motif of 4-1BB. 4-1BB binding to LANCL2 shows strong competition with GSH, albeit GSH binding affinity is nearly 20-fold weaker than of 4-1BB. Summary of results of nHU experiments carried out at increasing GSH concentrations. See supplemental Figure 9 for further details.

The utilization of a “Zinc-clasp” structural motif to stabilize signaling complexes is a powerful regulatory mechanism, notably exemplified by the coordination between CD4 and Lck in immune signaling ^49^ and between FEM1B and FNIP1 in metabolic regulation ^50^. Both TRAF2 and LANCL2 are Zn^2+^ ion binding proteins. Whereas in TRAF2, this site is located outside the motif-binding domain, in LANCL2 this ion directly forms the active site. To test if Zn^2+^ ions are important for the interaction, we carried out nHU titration experiments with GFP-TRAF2 and GFP-LANCL2 in the presence of increasing EDTA concentrations, which strongly chelates divalent cations in the cell extract (Figure 4C, Supplemental Figure 9A, Supplemental Data 2). TRAF2 binding was completely insensitive of the presence of bound Zn^2+^ ion, but LANCL2 binding showed strong dependence, indicating that 4-1BB binds at the active site of LANCL2, possibly via a ‘Zinc-clasp’ (Figure 4C).

We observed that the putative LANCL2-binding motif localizes to a GCE tripeptide segment that resembles a reversed glutathione (GSH), a pseudo ECG tripeptide, which is the natural ligand of LANCL2. To test a potential competition between these ligands, we performed nHU titration experiments between 4-1BB and GFP-LANCL2 in the presence of various amounts of reduced glutathione (Figure 4D, Supplemental Figure 9B, Supplemental Data 2). Clear competition was found between GSH and 4-1BB, which confirmed that both target the same molecular ‘Zinc-clasp’ interface on LANCL2. Moreover, the observed competition could also be fitted to a competitive binding model revealing that GSH binding is approximately 20-fold weaker than 4-1BB binding with an affinity of around 18 μM dissociation constant (Figure 4D, Supplemental Figure 9B).

### Redox-dependence of 4-1BB interactions and their impact on cellular redox homeostasis

The 4-1BB tail contains multiple cysteine residues that could potentially undergo oxidation and intramolecular cross-linking. This can likely happen in a biological context as 4-1BB activation is known to lead to increased reactive oxygen species (ROS) generation and thus a shift of cellular redox state to a more oxidative environment. This can also occur artificially in our in vitro nHU experiments. To prevent this, our cell lysis buffer contained 5 mM TCEP (tris(2-carboxyethyl)phosphine), a strong reducing agent ensuring reduced state during the nHU experiments. We tested how changes in the redox environment affect 4-1BB interactions by adding various amounts of TCEP and H2O2 to identical aliquots of cell extracts containing either GFP-TRAF2 and GFP-LANCL2 immediately prior to nHU titration experiments (Figure 5A, Supplemental Figure 10, Supplemental Data 2). By doing so, we established various ‘net reducing power’ in these binding experiments, which we defined as the molar concentration of H2O2 subtracted from the molar concentration of TCEP ([TCEP] – [H₂O₂]). Potentiometric measurements confirmed that the redox environment of our mixtures followed the expected dependence on the defined ‘net reducing power’ and that the presence of dilute cell extract did not substantially alter the measured oxidation–reduction potential. Mixtures remained in a reducing state until the externally added oxidant did not exceed the reducing capacity of TCEP. Around the point where the reducing capacity of TCEP was balanced by the added H₂O₂, the system shifted into the oxidizing range.

**Figure 5.**
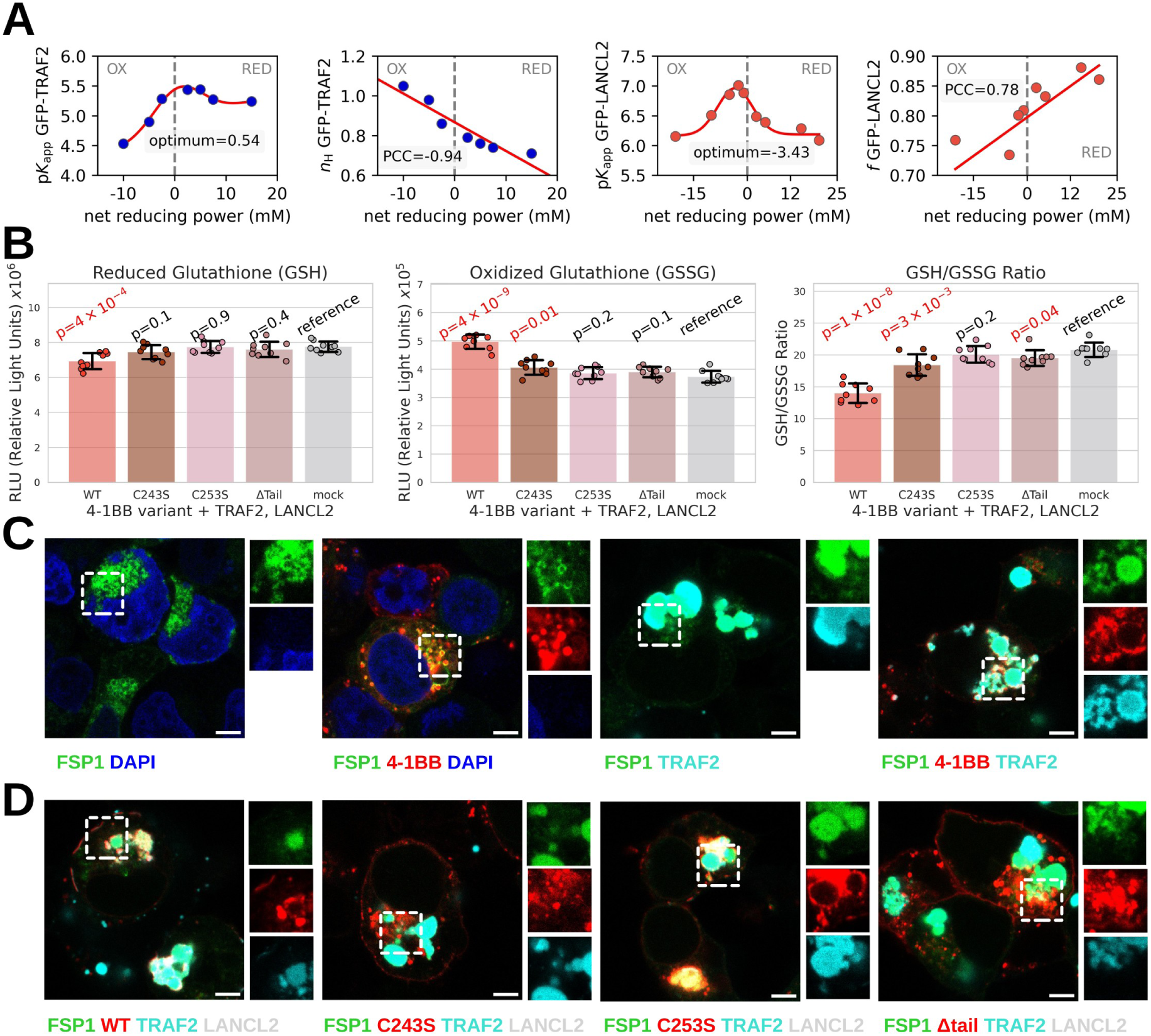
Redox-regulation and signaling of 4-1BB. (A) The binding between 4-1BB and TRAF2 (blue, on the top left) or 4-1BB and LANCL2 (red, on the top right) was measured in the presence of varying amounts of H2O2 and TCEP creating varying redox conditions. Both interactions show optimal binding at near physiological conditions with approximately an order of magnitude affinity difference depending on the redox state. TRAF2 shows optimal binding at slight excess of TCEP while LANCL2 binding prefers mildly oxidative environment. In addition, both interactions show changing binding models depending on redox condition. TRAF2 loses cooperativity and LANCL2 shows increased binding inactive fraction at increasingly oxidative conditions. (B) 4-1BB expression in HEK293T cells reduced GSH levels, increased GSSG levels, and decreased GSH/GSSG ratio. Tail variants had minimal effects, except for a modest oxidation shift observed for the C243S and Δtail. See supplemental Figure 10 for further details. (C) Fluorescence images of HEK293T cells transfected with FSP1, or combinations with 4-1BB and TRAF2. (D) Fluorescence images of different 4-1BB variants with FSP1, TRAF2 and untagged LANCL2. Confocal images were taken from fixed HEK293T cells transfected with fluorescently tagged proteins. Red, green, or cyan colors indicate mCherry, eGFP, or CFP tags, respectively. DAPI staining is shown in blue. For each panel, an overlay signal is shown on the left and the insets on the right show separate channels. Scale bar is equivalent to 5 μm. Representative images are shown based on hundreds of observations.

For both interactions, a clear binding optimum was identified under mild redox conditions. In the case of TRAF2, this experiment revealed that optimal binding occurs in a slightly reducing environment and that the cooperative nature of the interaction gradually disappears in oxidizing environment. In the case of LANCL2, the binding optimum was observed in mildly oxidizing environment, likely due to the oxidation of GSH into GSSG in the extract, and we also found that the partial binding activity of LANCL2 gradually increased in reducing environment, also likely due to GSH depletion (Figure 5A). These results clearly demonstrate that both interaction are redox-dependent and suggest that cellular redox fluctuations could dynamically modulate 4-1BB signaling.

To investigate whether the studied 4-1BB complexes influence cellular redox homeostasis, we measured intracellular GSH and GSSG levels in HEK293T cells expressing mCherry-tagged 4-1BB variants together with CFP-TRAF2 and GFP-LANCL2 (Figure 5B). We observed a marked shift toward an oxidative state, characterized by decreased GSH and increased GSSG levels, but only in the presence of WT 4-1BB together with TRAF2 and LANCL2. In contrast, the C243S mutant of 4-1BB, which disrupts TRAF2 interaction, induced only a slight oxidative shift compared to mock-transfected cells. Even smaller oxidative change was found with the C-terminal truncation of 4-1BB and the expression of the C253S mutant resulted in no detectable changes in redox balance (Figure 5B). Together, these results indicate that intact 4-1BB–TRAF2 complex formation is associated with a shift toward a more oxidative intracellular redox state, potentially promoting ferroptosis where LANCL2 interaction could occur due to the absence of the competitor GSH.

A recent study showed that the phase separation of ferroptosis suppressor protein-1 (FSP1) promotes ferroptosis ^51^. Thus, we examined if FSP1 localization could serve as a cellular marker of changes in cellular redox state upon expression of 4-1BB or TRAF2. Interestingly, FSP1 showed clear co-localization with both internalized 4-1BB-positive vesicles and TRAF2 signalosomes (Figure 5C). When 4-1BB, TRAF2, and FSP1 were co-expressed, the localization pattern of FSP1 closely resembled that of LANCL2 and removal of the complete 4-1BB tail abolished this interaction (Figure 5D). Notably, the co-localization of 4-1BB with FSP1 was markedly reduced upon introduction of the 4-1BB C243S mutation suggesting that FSP1 recruitment is mostly controlled by TRAF2 phase separation (Figure 5D).

### LANCL2 expression modulates downstream 4-1BB signaling

Besides redox regulation, one of the most characterized signaling output of 4-1BB activation is the NF-κB signaling. To investigate the contribution of 4-1BB, TRAF2, and LANCL2 on this downstream pathway, we established a NF-κB activity reporter HEK293T cell line, termed here as HEK293T-kb-luc, enabling the detection of NF-κB pathway activation through luminescence, and measured NF-κB activation upon transient expression of each protein individually, in pairwise combinations, or in triple combination (Figure 6, Supplemental Figure 11). Among single transfections, only TRAF2 increased NF-κB activity. In pairwise combinations, co-expression of TRAF2 with 4-1BB strongly increased the basal TRAF2 activity and co-expression of LANCL2 with either partner did not increase NF-κB activity. Strikingly, when 4-1BB and TRAF2 were also present, LANCL2 further enhanced the signal, with maximal NF-κB activity observed upon co-expression of all three proteins (Figure 6). Together, these findings suggest that LANCL2 modulates 4-1BB-mediated NF-κB signaling and may act in concert with TRAF2 in this pathway.

**Figure 6.**
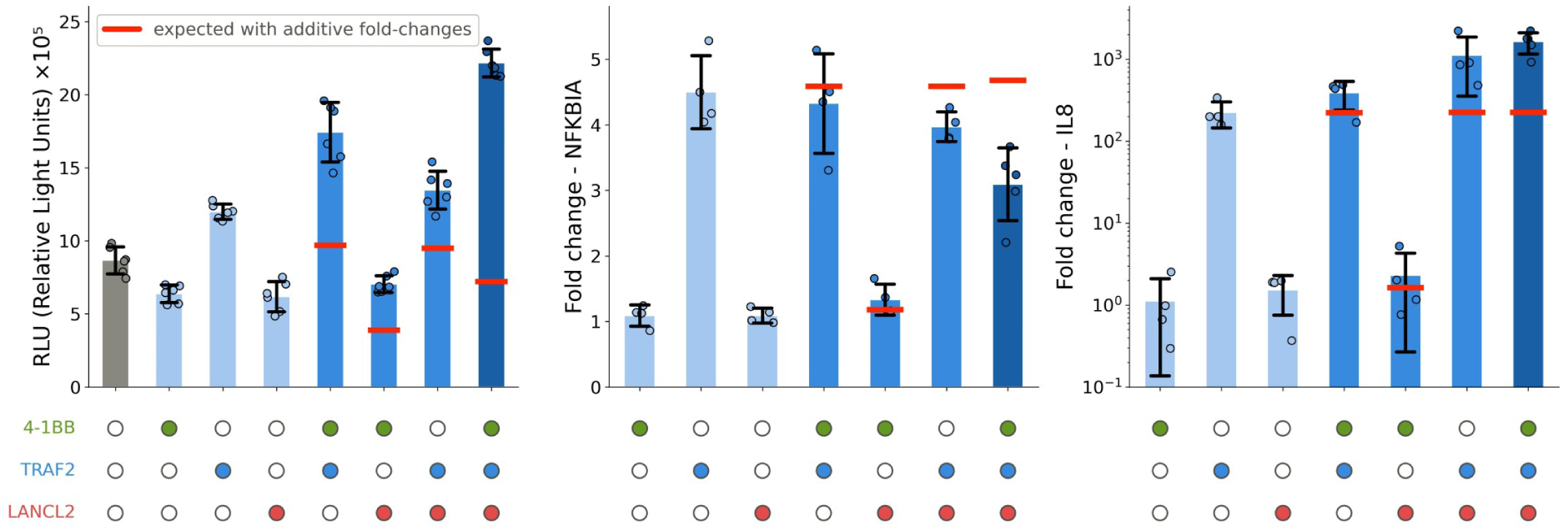
LANCL2 contributes to cellular NF-κB activity. (left) A HEK293T NF-κB luciferase reporter cell line was used to measure NF-κB activation after transfection with different combinations of 4-1BB, TRAF2, and LANCL2. Maximal NF-κB activity was observed when all partners were co-expressed. (middle) The expression of early response genes of NF-κB signaling, such as NFKBIA, are spontaneously driven by TRAF2 expression in HEK293T cells which is mildly suppressed in the presence of 4-1BB and LANCL2. (right) The expression of the early cytokine gene IL8 is not enhanced by the presence of LANCL2 alone, but is strongly enhanced when co-expressed with TRAF2. When TRAF2 is co-expressed with LANCL2 or 4-1BB and LANCL2 increased IL8 transcription was found. Fold change indicates 2^-ΔΔCt^ values to non-transfected cells using RPLP0 house keeping control. Expected levels assuming additive effects are shown with red horizontal dashed lines on each relevant bars. See supplemental Figure 11 for further details.

To examine these effects at the level of endogenous transcriptional responses, we measured the expression of known NF-κB target genes upon transient expression of 4-1BB, TRAF2, and LANCL2, either individually or their combinations in HEK293T cells by qRT-PCR (Figure 6). The early response genes *NFKBIA* and *TNFAIP3* ^52^ showed similar effects where TRAF2 expression alone strongly induced transcription, which was mildly attenuated in the presence of both LANCL2 and 4-1BB (Figure 6). The expression of *IL8*, a rapidly inducible NF-κB target cytokine gene ^52,53^, was also enhanced by TRAF2, whereas LANCL2 alone had no effect. In contrast, co-expression of TRAF2 and LANCL2 resulted in a marked increase in *IL8* expression (Figure 6). Together, these data show that LANCL2 modulates NF-κB-dependent transcriptional outputs and can potentiate TRAF2-driven signaling in a gene-specific manner.

## Discussion

### Old complex, new binding mechanism

The molecular connection between 4-1BB and TRAF2 was first reported in 1998 by three independent groups ^13,14,54^. Remarkably, just a year later, the crystal structure of the core complex was determined ^19^, and in 2000, the same group performed detailed calorimetric analyses of multiple TRAF2 complexes, among which 4-1BB unexpectedly exhibited little to no measurable binding ^20^. Despite these productive early efforts, research on this interaction soon lost momentum, and it effectively became a “cold case.”

By revisiting this long-standing question, we provide the first quantitative biophysical characterization of the 4-1BB–TRAF2 interaction, successfully determining a binding affinity within the typical range of motif-mediated interactions ^55^. Leveraging a next-generation interactomic tool called native holdup, or shortly nHU ^24^, we show that the binding affinity between TRAF2 and 4-1BB is over 400-fold stronger than previously measured (noting that the earlier measurement used a smaller mouse peptide) and precisely map the TRAF2 interaction site on 4-1BB. While confirming the importance of the acidic residues ^13^, our analysis also revealed a previously unrecognized requirement for a conserved cysteine: substitution with serine (C243S) abolished binding.

Analysis of human genetic variations of 4-1BB further supports the functional relevance of the refined TRAF-binding site. In the gnomAD database (v4.1.0) ^56^, no missense variants affecting cysteine 243 were identified, whereas the adjacent cysteines showed rare substitutions (C241Y, 87 alleles, no homozygotes; C253S, 20 alleles, including one homozygote; C253Y, 2 alleles, no homozygotes; no phenotype data are available for any of these observed variants). The absence of variants at the TRAF-binding cysteine suggests strong evolutionary constraint, consistent with its essential role in 4-1BB signal transduction. In line with this, we identified several other partners of TRAF proteins that contain Cys residues at an analogous position, such as in CD27.

### Implications on 4-1BB signaling

Our findings refine the current model of 4-1BB signaling by positioning the receptor as a redox-responsive regulatory hub rather than a purely TRAF-driven activator. Beyond TRAF2, our data reveal that the 4-1BB cytoplasmic tail can also engage two redox-related proteins, LANCL2 and GPX1. GPX1 was previously identified as a binding partner of TRAF2 ^31^ and confocal images revealed that LANCL2 can co-localize with TRAF2 in cells, effectively highlighting this as a cognate signaling network. Our findings also suggest that 4-1BB may form distinct complexes depending on the cellular redox state.

Previous studies showed that 4-1BB costimulation is associated with enhanced mitochondrial activity and increased intracellular GSH and GPX4 expression, providing the proper metabolic fitness for clonal expansion of CD8+ T-cells ^57^. Consistent with this, GPX4 has been shown to protect T cells from the accumulation of membrane lipid peroxides and ferroptosis ^58,59^. Conversely, 4-1BB ligand overexpression has been reported to promote ROS production leading to decreased GSH levels and decreased GPX4 expression while its knockdown had the reverse effect in a mouse model ^60^. GPX7 expression was also found to deregulate TRAF2 stability ^61^. Together, these observations suggest that 4-1BB signaling may operate within a finely balanced redox window, in which metabolic activation is coupled to antioxidant defenses, while excessive ROS accumulation may shift the system toward lipid peroxidation and ferroptosis.

We now provide mechanistic support for this redox dimension of 4-1BB signaling by linking three redox-sensitive proteins to 4-1BB signaling: TRAF2, LANCL2, and GPX1. We redefine the TRAF2-binding region and show that although TRAF2 associates optimally under reducing conditions, mild oxidation promotes site-specific disulfide formation that stabilizes the TRAF2–4-1BB complex. This oxidative “TRAF2-lock” may prolong receptor engagement and is poised to amplify downstream signaling output, as well as adjacent mitochondrial ROS production. GPX1 appears to associate with 4-1BB independently of LANCL2, most likely indirectly via TRAF2, and likely provides local H₂O₂ detoxification, but only when GSH is sufficiently available, consistent with observations that early 4-1BB signaling increases intracellular GSH levels and supports GPX-dependent antioxidant defense.

As redox conditions shift from reduction to mild then moderate oxidation, the 4-1BB signaling machinery transitions into a distinct state defined by reinforced oxidative pressure and reconfigured protein interactions. The TRAF2 disulfide lock establishes a positive redox feedback loop: moderate ROS stabilizes TRAF2 on the receptor, which further sustains TRAF2-dependent mitochondrial ROS production. As oxidative load increases, local GSH becomes limiting, leading to progressive loss of GPX1 and GPX4 activity consistent with the results of 4-1BB ligand overexpression experiments. Under these conditions, the active site of LANCL2 becomes gradually liberated from GSH and its ‘Zinc-clasp’ interaction with 4-1BB becomes the favored, which was out-competed in reducing stage by GSH due to its large cellular concentration, despite having 20-fold weaker affinity for LANCL2 than 4-1BB. This oxidative reconfiguration of the 4-1BB interactome provides a mechanistic framework linking 4-1BB signaling to ferroptotic sensitivity: prolonged TRAF2-driven ROS production, declining GSH, weakened GPX activity, and increased LANCL2 engagement collectively shift the cell toward a redox state associated with lipid peroxidation and ferroptosis, which is also supported by the cellular co-localization with FSP1. In addition, LANCL2 expression was already found correlated with expression of ferroptosis marker proteins ^62^ and decreased GPX1 levels increases susceptibility to ferroptosis ^63^. Together, these findings integrate previously disconnected observations and reveal 4-1BB as a sensor and interpreter of redox cues, switching between antioxidant-supported signaling and pro-oxidative, ferroptosis-sensitizing signaling depending on the cellular redox state (Figure 7).

**Figure 7.**
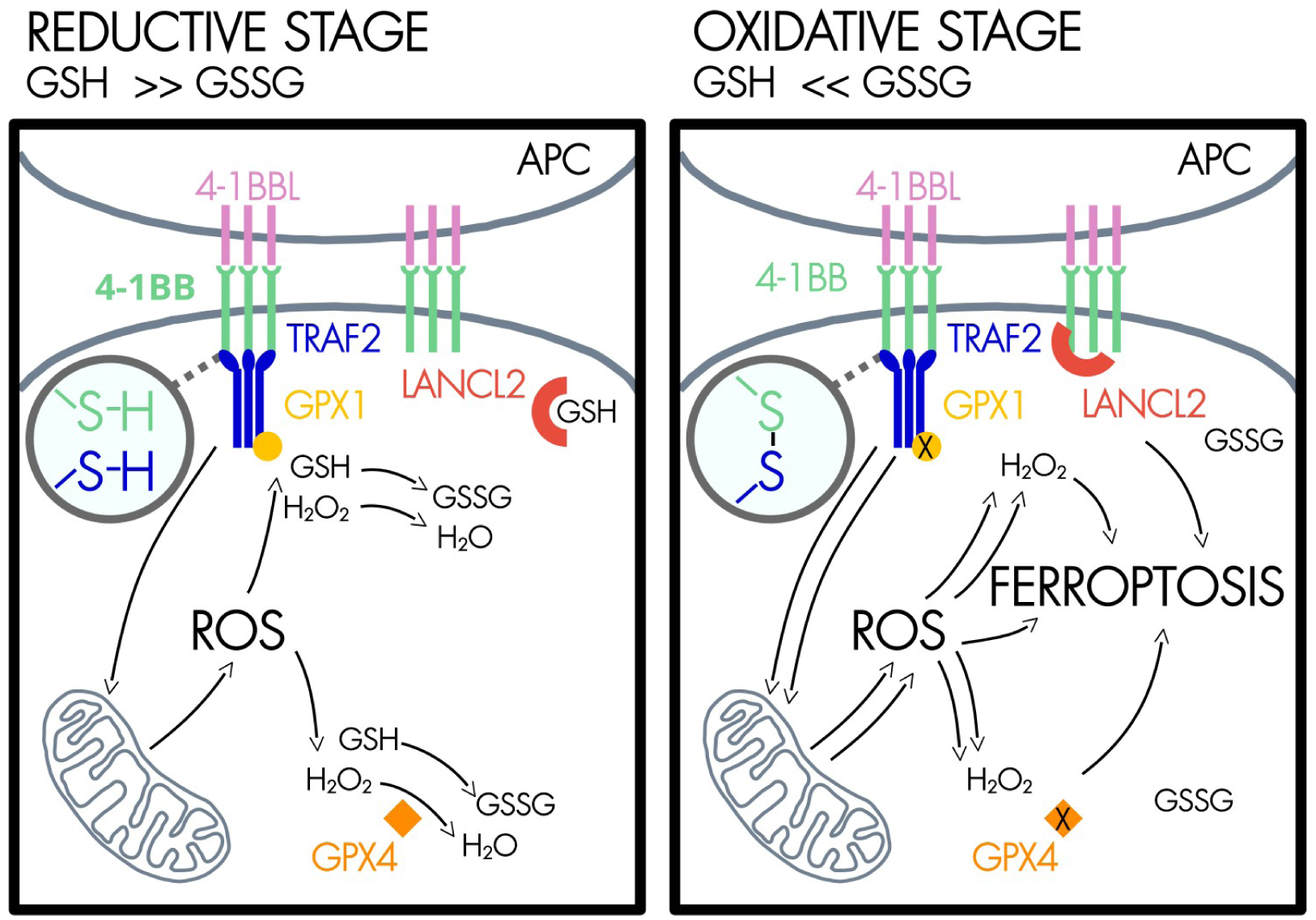
4-1BB signaling regulates cellular redox homeostasis. 4-1BB activation generates ROS leading to a shift towards a more oxidizing intracellular environment. This redox change modulates 4-1BB interactions. TRAF2 interaction may be stabilized via a disulfide bond. Stronger oxidation reduces cellular GSH levels, allowing LANCL2 binding. Together, these events may contribute to oxidative stress and may promote ferroptosis.

Beyond redox regulation itself, our NF-κB reporter and transcriptional analyses suggest that changes in the composition of the 4-1BB signaling complex are accompanied by changes in downstream signaling output. Although LANCL2 alone had limited effects on NF-κB activity, its co-expression with TRAF2 and 4-1BB clearly modulated NF-κB-dependent signaling. Notably, the effects on the expression profile of endogenous NF-κB target genes were gene-dependent, indicating that LANCL2 does not simply amplify canonical NF-κB activation but may influence the qualitative transcriptional output of the pathway. This observation is consistent with a model in which redox-dependent remodeling of the 4-1BB interactome regulates not only signaling magnitude but also signaling specificity.

### Implications for cancer therapy and immune modulation

Our findings have implications for inflammatory disease, cancer immunotherapy, and engineered cell therapies. First, the observation that omilancor, an oral LANCL2 pathway activator currently in phase 3 clinical trials for ulcerative colitis and phase 2 for Crohn’s disease, enhances oxidative metabolism in regulatory T cells ^64^ gains more weight in light of the interaction with 4-1BB and their link to redox signaling. Second, the fact that LANCL2 binding is centered on Cys253 in 4-1BB that is mutated to Tyr in the mouse sequence reveals a human-specific component of 4-1BB signaling. This limitation possibly affects the pre-clinical assessment of 4-1BB agonists and toxicity in cancer immunotherapy pipelines which are typically optimized and validated in murine systems lacking this interaction. Finally, the redox-dependent stabilization of TRAF2 and the local recruitment of GPX1 and LANCL2 demonstrate that the 4-1BB cytoplasmic tail operates as a redox-responsive signaling hub rather than a simple TRAF2-binding ‘costimulatory module’ in CAR constructs. These findings therefore suggest that the efficacy and activity of 4-1BB-based therapies may be influenced not only by receptor engagement, but also by the redox and metabolic state of the responding cell and by human-specific protein interactions. Incorporating these variables into the design and pre-clinical evaluation of 4-1BB agonists and engineered immune-cell therapies may be important for achieving more predictable immune activation while limiting unwanted toxicity.

## Author contributions

BZ, BJ, EK performed all cloning and extract preparation for nHU experiments. GG, BZ, BJ, EK performed all nHU experiments and their analysis. SK, AT, DMP, BM, LN, MV performed mass spectrometry measurements and their analysis. CB, AZ performed data analysis. BZ conceptualized and performed the cellular functional assays. BJ, ALW, GS performed the SimPull experiments and their analysis. GG and MV conceived and supervised the project; GG wrote the original draft and all authors refined the paper.

## Competing interests

The authors declare no competing interests.

## Acknowledgement

We thank Aurelie Rossin and Anne-Odile Hueber for providing the 4-1BB template for cloning. We are grateful to Camille Kostmann for her help in the creation of the Sleeping Beauty gene delivery system with the puromycin resistance cassette. The authors would like to thank Sameh BEN AICHA and Mathis COURTILLET, engineers at the PRISM microscopy platform at the Valrose Institute of Biology, which forms part of the ‘Microscopy, Imaging and Cytometry Azur’ platform accredited by the GIS – IbiSA, for their help in microscopy. G.G. was supported by Inserm, Université Côte d’Azur (IdEx Attractivity Package, CSI, MSI-MIDI), Fondation ARC pour la recherche sur le cancer (PJA1, 2024), the ATIP-Avenir program, and the T-ERC STG 2024 grant (ANR-24-ERCS-0002). This work is part of the “Orphan Motifs” project led by GG, supported by the Impulscience program of the Fondation Bettencourt Schueller. M. V. received support from Research Council of Finland (nos. 288475 and 294173), the Sigrid Jusélius Foundation, the Finnish Cancer Foundation, Biocenter Finland, and HiLIFE. B.Z. was supported by CNRS, and by the Université Côte d’Azur (IdEx Attractivity Package ANR-15-IDEX-01).

## Methods

### Peptide synthesis

The C-terminal tail peptide of 4-1BB including residues 230-255, as well as all its mutant versions were obtained commercially from Synpeptide or GenicBio with an N-terminal biotin label attached to the N-terminus via an Ahx (6-Aminohexanoic Acid) linker and with free C-terminus. The CD27 peptide including residues 214-260 was obtained commercially from GenicBio with an N-terminal biotin label attached to the N-terminus via an Ahx linker and with free C-terminus. All peptides were HPLC purified to >95% purity. Predicted peptide mass was confirmed by MS and peptide concentration was determined based on dry weight.

### Cloning, mutagenesis

The full-length sequences of LANCL2, TRAF2, TRAF5 and PARD3 were obtained from cDNA pools and were cloned into pEGFP-C1 or pECFP-C1 vectors with N-terminal GFP or CFP tag, respectively. A GS linker was introduced between the fluorescent tag and protein of interest. Fsp1/AIFM2 was cloned into pEGFP-N1 plasmid for C-terminal GFP tagging. Sequences were verified by Sanger or nanopore sequencing (Eurofins Genomics Germany GmbH, Ebersberg, Germany). The full-length sequence of 4-1BB was cloned into a pCEP4-mCherry mammalian expression vector with C-terminal mCherry tag. TRAF2 and 4-1BB mutations were introduced by standard site-directed mutagenesis (QuickChange) strategy using Platinum SuperFi II polymerase (Thermo Fisher Scientific, Bremen, Germany), and the mutations were confirmed by sequencing. The last 9 residues of 4-1BB were removed to obtain the Δtail 4-1BB construct.

### Extract preparation and manipulation

All cells were confirmed to be mycoplasma-negative prior to use using the Eurofins Mycoplasmacheck qPCR-based service.

Jurkat E6.1 cells (ECACC #88042803, RRID: CVCL_0367, not authenticated) were grown in RPMI (Gibco) medium completed with 10% FBS (Gibco BRL) and 40 μg/ml gentamicin (Gibco/Life Technology). To prepare total cell extracts, Jurkat cells were seeded onto T-175 flasks and grew until 3×10^6^ cells/ml confluency. Cells were collected by 1,000 g x 5 min centrifugation, washed with PBS and then lysed in ice-cold lysis buffer (Hepes-KOH pH 7.5 50 mM, NaCl 150 mM, Triton X-100 1%, cOmplete EDTA-free protease inhibitor cocktail 5x, TCEP 5 mM, glycerol 10%).

HEK293T cells were authenticated by short tandem repeat (STR) profiling using ATCC Cell Line Authentication Service (Manassas, VA, USA), showing a 100% match to ATCC ref. CRL-3216 (RRID:CVCL_0063, the authentication report is available upon request). The HEK293T cells were grown in Dulbecco’s modified Eagle’s medium (DMEM, Gibco, glucose: 4.5 g/liter, 110 mg/liter sodium pyruvate) completed with 10% FCS and 1% Penicillin-Streptomycin (Gibco) on CellBIND flasks (Corning), diluted 1:10 every ∼2nd/3rd day. All cells were kept at 37°C and 5% CO 2. For transient transfection prior to lysate preparation, cells were seeded on poly-D-lysine (Sigma-Aldrich, ref. P6407) treated 100 mm cell culture dishes (2×10^6^ cells/dish). The next day, the transfection was carried out with JetPRIME reagent (Polyplus, France) according to the manufacturer’s recommendations using 5 μg total DNA and 20 μl of JetPRIME reagent/well. The medium was replaced with fresh medium after 5 hours of treatment with the transfection mixture. Two days after transfection, cells were washed with PBS and then lysed in ice-cold lysis buffer (Hepes-KOH pH 7.5 50 mM, NaCl 150 mM, Triton X-100 1%, cOmplete EDTA-free protease inhibitor cocktail 5x, TCEP 5 mM, glycerol 10%).

Cell suspensions were sonicated 4x20 sec with 1 sec long pulse on ice, then incubated rotating at 4°C for 30 min. The lysates were clarified by centrifugation at 12,000 rpm 4°C for 20 min and supernatant was kept. Total protein concentration was measured by standard Bradford method (Bio-Rad Protein Assay Dye Reagent #5000006) using a BSA calibration curve (MP Biomedicals #160069, diluted in lysis buffer) on an Eppendorf BioPhotometer spectrophotometer or on a Thermo Scientific NanoDrop One instrument in cuvette mode. Lysates were diluted to 2 mg/ml concentration, aliquoted and snap-frozen in liquid nitrogen and stored at −80°C until measurement.

Fluorescence levels in extracts were measured before snap-freezing or immediately prior to the experiment to determine the optimal dilution, ensuring a signal at least 20-fold above background. Measurements were performed in 384-well low-binding microplates (E18063G5, Greiner Bio-One, Kremsmünster, Austria) using a SpectraMax iD5 multimode plate reader (Molecular Devices) with an excitation wavelength of 485 ± 15 nm and an emission wavelength of 530 ± 20 nm for GFP and with an excitation wavelength of 410 ± 15 nm and an emission wavelength of 465 ± 20 nm for CFP. Final dilutions to working concentrations (2 mg/ml for mass spectrometry and 0.1-1mg/ml for fluorescence-based nHU experiments depending on fluorescence level) were made on the thawed samples right before the experiment using corresponding lysis buffer.

In certain experiments, the freshly thawed cell extracts were mixed with varying amounts of EDTA, reduced glutathione, hydrogen peroxide, or TCEP. For the changing redox conditions, the redox potential of free mixtures of hydrogen peroxide and TCEP in lysis buffer, or the mixtures of hydrogen peroxide and TCEP in lysis buffer containing 0.1-1 mg/ml total cell extract was measured using an InLab Redox Micro electrode connected to a FiveEasy Plus device (Mettler Toledo).

### Single point and titration nHU experiments

Single-point nHU experiments were carried out at ∼10 μM empirically estimated bait concentration. For this, 50 μl streptavidin resin (Streptavidin Sepharose High Performance, Cytiva) was incubated with 250 μl 70 μM biotinylated peptides (or biotin for controls) in 0.5 ml Protein LoBind tubes (Eppendorf). After mixing, reaction mixtures were incubated overnight at 4 °C, or for 1 hour at room temperature. After the incubation, all resins were washed with 9 resin volume (450 μl) holdup buffer (50 mM Tris pH 7.5, 300 mM NaCl, 1 mM TCEP, 0.22 filtered). The washed resins were then mixed with 25 μl 1 mM biotin solution, diluted in 10 resin volume holdup buffer and were incubated for 3 minutes at room temperature. Then, the resulting resins were washed four times with 9 resin volume holdup buffer. The resulting bait-saturated resins were mixed with 200 μl 2 mg/ml Jurkat extracts and were incubated at 4 °C for 2 h with constant mild agitation. After the incubation ended, the solid and liquid phases were separated by a brief centrifugation (15 sec, 2000 g) and 130 μl of the supernatant was recovered rapidly. Then, to minimize carryover contamination from resin, the recovered supernatants were centrifuged one more time and 100 μl of the supernatant was recovered that was subjected for mass spectrometry analysis.

Titration holdup experiments were carried out as described above using 25 μl saturated resins prepared. Briefly, we mixed control, or bait-saturated resins at various proportions by keeping the total resin amount and resin-analyte ratio constant during the experiment (for 25 μl we used 100 μl cell extract as analyte).

Experiments were carried out either in single tubes, as described above, or on filter plates with identical results. For on-plate single point measurements, resin saturation was carried out directly on Pall (Acropep Advance 350 1.2 Supor) filter plates with 25 μl resin volume. The plate containing resin and peptides or biotin control was mixed every 10 minutes for 1 hour at room temperature at 800 rpm on an Eppendorf MixMate. The plates were incubated ON at 4 °C. All washes were carried out using a vacuum manifold or by centrifugation. The prepared plate with bait-saturated resins were mixed with 100 μl 2 mg/ml cell extracts and were incubated at 4 °C for 2 h with constant mixing with 700 rpm on a BioSan Mini-Shaker PSU-2T. For fluorescent measurements, more diluted cell extracts were used ensuring that at least 20-fold background fluorescence was achieved. Depleted extracts were collected on an LoBind deep well plate (Eppendorf, 1ml). For on-plate titration experiments, saturated resin stocks were prepared in tubes and resin mixing was done on filter plates and all additional steps were identical to the titration experiments.

### Liquid Chromatography–Mass Spectrometry analysis of nHU experiments

With the exception of Experiment #3 of the WT 4-1BB tail, all experiments were carried out on a Bruker timsTOF Pro2 instrument with data independent acquisition (DIA). The 100 µl nHU samples were TCA precipitated by adding 20 µl 100% TCA (#100807, Merck). Immediately after TCA addition the samples were mixed well with vortex and left overnight to 4 °C to precipitate. The following day the samples were centrifuged at 13 000 rpm for 15 min at 4 °C. The protein pellets were washed with 400 µl ice-cold 100% acetone, and centrifuged 13 000 rpm for 8 min at 4 °C. The supernatant was discarded and the pellets washed again with 400 µl ice-cold 100% acetone followed by centrifugation at 13 000 rpm for 5 min at 4 °C. After centrifugation the supernatant was carefully removed by pipetting and pellets were air dried. To drive off acetone the pellets were placed in 95 °C heat block for 5-10 min until no liquid was visible. 50 µl 8M Urea was added to the pellets, mixed with vortexing and put to −20 °C to solubilize. The following day the samples were vortexed and left overnight to 4 °C to solubilize. After the incubation 8M Urea was diluted with 100 mM Tris-HCl, pH 8.0 to 4M by adding 50 µl buffer, mixing by vortexing. Proteins were reduced with 5 mM (11 µl of 50 mM stock) TCEP for 30 min at 37 °C, and alkylated with 10 mM iodoacetamide (12 µl of 100 mM stock) for 30 min in the dark at room temperature. Urea concentration was diluted to ∼1 M by adding 300 µl 100 mM Tris-HCl, pH 8.0. The pH of the samples was controlled and 1 µg Trypsin/Lys-C Mix (V5073, c=0.5 µg/µl, Promega) was added to each sample and mixed by vortexing. Samples were digested overnight at 37 °C. Peptide mixtures were then desalted on a C18 mini columns (#HUM S18V, Higgins Analytical, Inc.), and dried in a centrifuge concentrator (Concentrator Plus, Eppendorf).

For MS acquisition the dried peptides were resuspended in 40 µl Buffer A (0.1% (vol/vol) TFA, 1% (vol/vol) acetonitrile (#83640.320, VWR) in HPLC grade water (#10505904, Fisher Scientific). For the DIA analysis, the resuspended peptides were further diluted 1:20 in buffer A1 (1% formic acid in HPLC water). 20 µl was loaded into an EvotipPure (Evosep Biosystems, Denmark) according to the manufacturer’s instructions and run at the same time as three technical replicates.

The samples were analyzed using the Evosep One liquid chromatography system coupled to a hybrid trapped ion mobility quadrupole TOF mass spectrometer (Bruker timsTOF Pro2, Bruker Daltonics) ^65^ via a CaptiveSpray nano-electrospray ion source (Bruker Daltonics). An 8 cm × 150 µm column with 1.5 µm C18 beads (EV1109, Evosep) was used for peptide separation with the 60 samples per day method (21 min gradient time). Mobile phases A and B were 0.1 % formic acid in water and 0.1 % formic acid in acetonitrile, respectively. The MS analysis was performed in the positive-ion mode with dia-PASEF method ^66^ with sample optimized data independent analysis (dia) scan parameters. We performed DDA in PASEF mode from a pooled sample to be able to adjust dia-PASEF parameters optimally. To perform sample-specific dia-PASEF parameter adjustment, the default dia-short-gradient acquisition methods were adjusted based on the sample-specific DDA-PASEF run with the software “tims Control” (Bruker Daltonics). The ion mobility windows were set to best match the ion cloud density from the sample type-specific DDA-runs. The following parameters were used in dia-PASEF runs: TIMS settings 1/KO 0.85-1.30, Ramp time 100 ms, Accumulation time 100 ms, Duty cycle 100%, and dia-PASEF settings cycle time estimate 1.17 s, MS1 ramps 1, MS/MS ramps 10, MS/MS windows 18, mass range 357.2-1257.2 Da, Mobility range 0.85-1.30 1/K0. For collision energy at 0.60 1/KO 20 eV and at 1.60 1/KO 59 eV were used.

To analyze diaPASEF data, the raw data (.d) were processed with DIA-NN v1.8.1 ^67,68^ utilizing spectral library generated from the UniProt human proteome (UP000005640, downloaded 05.04.2022 as a FASTA file, 20358 proteins). During library generation, the following settings were used with fixed modifications: carbamidomethyl (C); variable modifications: acetyl (protein N-term), oxidation (M); enzyme:Trypsin/P; maximum missed cleavages: 1; mass accuracy fixed to 1.5e-05 (MS2) and 1.5e-05 (MS1); Fragment m/z set to 100-1700; peptide length set to 7-30; precursor m/z set to 300-1800; Precursor charges set to 2-4; protein inference not performed. All other settings were left to default.

Experiment #3 was carried out using an Orbitrap Exploris 480 with data dependent acquisition (DDA). Protein mixtures were TCA-precipitated overnight at 4°C, pellets were washed twice with 1 mL cold acetone, dried and dissolved in 2 M urea in 0.1 M Tris-HCl pH 8.5 for reduction (5 mM TCEP, 30 min.), and alkylation (10 mM iodoacetamide, 30 min.). Trypsin digestion was carried out at 37°C and overnight. Peptide mixtures were then desalted on C18 spin-column and dried on Speed-Vacuum before LC-MS/MS analysis. Samples were resuspended in 0.1% TFA and then analyzed using an Ultimate 3000 nano-RSLC (Thermo Scientific, San Jose California) coupled in line with a quadrupole - orbitrap Exploris 480 via a nano-electrospray ionization source (Thermo Scientific, San Jose California) and the FAIMS pro interface. Less than 200 ng tryptic peptides were loaded on a C18 Acclaim PepMap100 pre-column (300 µm ID x 5 mm, 5 µm, 100Å, Thermo Fisher Scientific) for 1 minutes at 15 µL/min with 2% ACN, 0.1% FA in H_2_O and then separated on a C18 PepMap nano-column (75 µm ID x 25 cm, 2.6 µm, 150Å, Thermo Fisher Scientific) with a 70 minutes linear gradient from 7% to 35% buffer B (A: 0.1% FA in H_2_O; B: 0.1% FA in 80% ACN) followed by a regeneration step at 90% B and a equilibration at 7% B. The total chromatography was 95 minutes at 450 nl/min and the oven set to 45°C. The mass spectrometer was operated in positive ionization mode in Data-Dependent Acquisition (DDA) for two FAIMS compensation voltages (CV = −45V & −55V). The DDA cycle consisted of one survey scan or MS1 (330-1400 m/z, 90,000 FWHM) followed by MS2 spectra (HCD; 30% normalized energy; 8 m/z window; Scan Range 140-1400 m/z; 22,500 FWHM). The cycle time was limited at 1.5 sec for the CV −45 and 1.2 sec for CV −55. The Automatic Gain Control (AGC target) were set to 3E6 and 1E5 respectively for MS1 and MS2 scans, and the maximum injection time (IT) was set to 50 ms for both scan modes. Unassigned and single charged states were rejected. The exclusion duration was set for 30s with mass width was ± 10 ppm.

Proteins were identified with Proteome Discoverer 2.5 software (Thermo Fisher Scientific) and *Homo Sapiens* proteome database (Swissprot, reviewed, release 2024_02_07 with 20598 entries). Precursor and fragment mass tolerances were set at 10 ppm and 0.02 Da respectively, and up to 2 missed cleavages were allowed. Oxidation (M) and Acetyl (Protein N-term) was set as variable modification, and Carbamidomethylation (C) as fixed modification. Peptides were filtered with a false discovery rate (FDR) at 1%, rank 1. Proteins were quantified with a minimum of 1 unique peptide based on the XIC (sum of the Extracted Ion Chromatogram).

### Mass spectrometry data processing

The measured extracted ion chromatogram (XIC) intensities or the quantified protein intensities from the DIA-NN Report.pg_matrix were pre-processed in Perseus. The measured protein intensities were normalized based on median intensities of the entire dataset to correct minor loading differences. For statistical tests and enrichment calculations, not detectable intensity values were treated with an imputation method, where the missing values were replaced by random values similar to the 10% of the lowest intensity values present in the entire dataset. These Perseus processed files were further refined. First, the normalization described above assumes that there is no major change in total protein content. To check this, we fitted the obtained log _2_ fold-change histogram to a normal distribution to determine the mean value and the width (2 σ) of the distribution of the majority of the proteome that do not show specific binding. Then, we tested whether the expected mean of 0 is within the 95% confidence interval of the measured mean. In case it was outside, we applied the mean as a correction factor for re-normalization to correct systematic change in protein content. Unpaired two tailed T-test, assuming equal variance, were performed on obtained log_2_ intensities. For each experiments, a hyperbolic significance threshold was applied as described before ^24^. This was calculated as

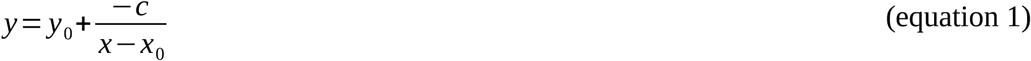

where *y* is the *P* value threshold at fold change of *x*, *c* is a curvature parameter empirically fixed at 2, *y*_0_ is the minimal *P* value threshold that a significant interaction needs to achieve (defined as 1.3 −log_10_P), and *x*_0_ is the width (2 σ) of the fold-change distribution of all detected proteins, calculated by fitting histograms with a Gaussian curve.

Obtained fold-change values were converted to fractional depletion values, *θ* as:

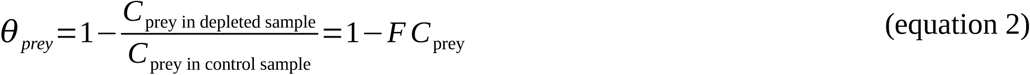

where *C* indicates the concentrations of the prey molecule and FC is the measured fold-change of intensity. Obtained degree of binding values were converted to apparent affinities using the hyperbolic binding equation assuming bimolecular complexes and ideal binding:

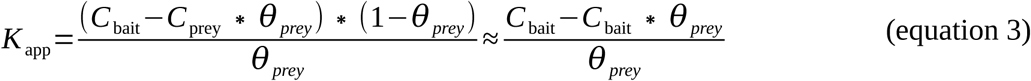

where *θ* denotes the degree of binding, *C*_prey_ and *C*_bait_ are the prey and bait molecule concentrations and *K*app is the apparent dissociation constant.

Since degree of binding *θ* values are intrinsic properties of each interactions under the same conditions, they are possible to combine and minimize the inaccuracy of each experiment. To get a smoothed data for the WT 4-1BB interactome, we took the average of *θ* from independent datasets for each partner. To pair an updated P-value for each protein, we used the Wilkinson’s method ^69^. Simply, we took into consideration the measured P-values of the experiment where the interaction was found to be the least significant and the number of experiment series (N) as

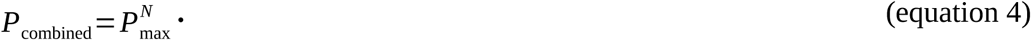

To estimate absolute concentrations and binding stoichiometries from nHU experiments, we used the obtained intensity values of the control samples and assumed that the total measured intensity is proportional and representative of the 200 mg/ml total cellular protein content ^70^. Then, we used the measured degree of binding values and combined these resources to obtain estimated bound concentrations of preys in the assay. As of note, these calculations are rather only order of magnitude estimations than precise concentration measurements. Nevertheless, by calculating the bound GPX1/TRAF2 ratios from the two experimental datasets, we obtained comparable stoichiometries with an average value of 1:6.6 with a standard deviation of 1.7 (Supplemental Data 1).

### Western blot and fluorescence analysis of nHU experiments

For Western blot, samples were mixed with 4x Laemmli Sample Buffer in 3:1 ratio. Equal amounts of samples were loaded on 10-12% acrylamide-gels. Transfer was done into PVDF membranes using a Trans-Blot Turbo Transfer System and Trans-Blot Turbo RTA Transfer Kit (BioRad, #1704275). After 1 hour of blocking in 5% milk, membranes were incubated overnight at 4°C with primary antibodies in 5% milk-TBS-Tween. The following antibodies and dilutions were used: TRAF2 (clone F-2, Santa Cruz Biotechnology, sc-136999, 1:1000, RRID:AB_2209298); LANCL2 (Thermo Fisher Scientific, PA5-54063, 1:1000, RRID:AB_2643271); GPX1 (Proteintech, 29329-1-AP, 1:1000, RRID:AB_2918283), α-tubulin (clone DM1A, Invitrogen, 14-4502-82, 1:1000, RRID:AB_1210456), GAPDH (clone 6C5, Invitrogen, AM4300, 1:20,000, RRID:AB_437392). Next day, the membranes were washed three times with TBS-Tween and incubated at RT for 1 h in secondary antibody (Jackson ImmunoResearch, goat anti-rabbit(H+L) #111-035-003 RRID: AB_2313567 or goat anti-mouse(H+L) #115-035-003 RRID: AB_10015289) in 5% milk-TBS-Tween (dilution: 1:10,000). After washing three times with TBS-Tween, membranes were exposed to chemiluminescent HRP substrate (Pierce ECL Western Blotting Substrate #32209 or SuperSignal West Femto Substrate #34094, Thermo Fisher Scientific) and captured in a docking system (Amersham Imager 600, GE). In case of reprobing the membranes with a different antibody, they were either cut before the first antibody incubation or kept intact and re-exposed to other primary antibody raised in a different species. In the latter case, membranes were treated with 15% H_2_O_2_ for 30 minutes to remove secondary signal and reblocked before continuing the incubation with the second primary antibody. Densitometry was carried out on raw Tif images by using Fiji ImageJ 1.54p.

For fluorescence measurements, depleted samples collected on receiver plates or in individual tubes were transferred to 384-well micro-plates (low binding microplate, 384 well, E18063G5, Greiner Bio-One, Kremsmünster, Austria) in triplicates. Readout was carried out in a Molecular Devices SpectraMax iD5 multimode plate reader with an excitation wavelength of 485 ± 15 nm and an emission wavelength of 530 ± 20 nm for GFP and with an excitation wavelength of 410 ± 15 nm and an emission wavelength of 465 ± 20 nm for CFP.

### Binding curve fitting

Data fitting of titration experiments were performed in QtiPlot and data visualization was done using Matplotlib. Optimal binding was determined based on fitting and anything else than simple binding mechanisms was only considered in cases where the data well justified more complex binding mechanism. For simple binding, hyperbolic model was used as:

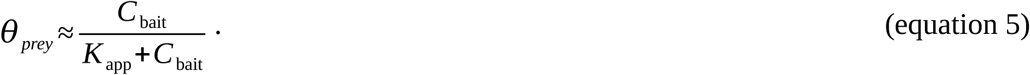

For interaction with LANCL2, optimal binding was observed with assumed partial binding:

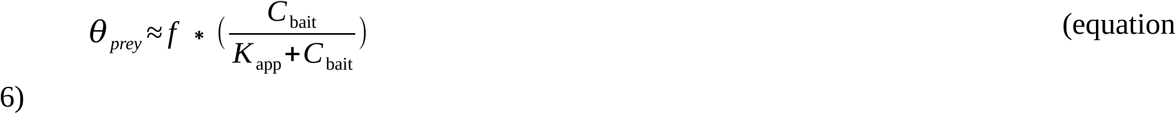

where the *f* factor is the “fraction of binding capable prey”. For TRAF2, negative cooperativity was observed and the titration curve was fitted to:

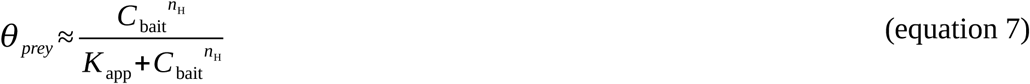

where the *n*H is the Hill coefficient. In the case when n_H_ is not equals 1, hence cooperativity is detected, *K*_app_ is an apparent binding affinity where

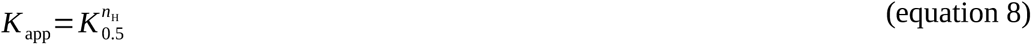

in which *K*_0.5_ is the ligand concentration producing half-maximal binding.

Endogenous TRAF2 bound with simple 1:1 behavior (*K*_app_ 2.5 µM), while GFP-TRAF2 showed slight negative cooperativity (*K*_app_ 2.7 µM). The comparable affinities indicate the GFP tag does not fundamentally alter the interaction; the difference in cooperativity may reflect the tag itself or the oligomerization state of overexpressed TRAF2. LANCL2 displayed partial binding activity, possibly reflecting occupancy of its active site by endogenous GSH in the cell extract, consistent with the competition between GSH and 4-1BB observed below (Figure 4D).

The analytical cubic competitive binding equation described in Equation 17 of Roehrl et al ^71^. was used to determine the affinity of GSH for LANCL2, using fixed parameters obtained from fitting the GSH-free titrations. Affinity dependence on EDTA or redox conditions was modeled assuming linear behavior. To determine the optimal redox condition, data were fitted with an empirical asymmetric Gaussian function allowing distinct plateaus at low and high redox values. Although simplified, this approach captures the overall macroscopic behavior of the system and robustly reflects the presence or absence of correlation.

### Immunostaining and Confocal microscopy

The day before transfection, 0.75×10^5^ HEK293T cells were seeded in the wells of a 24-well plate with coverslips. Cells were co-transfected with 0.5 µg of total plasmid DNA per well, with equal amounts of each plasmid in co-transfections (e.g., 0.25 µg of each plasmid for two-plasmid conditions), using JetPRIME reagent (Polyplus, #101000046) according to the manufacturer’s recommendations. The medium was changed to fresh medium after 5 hr of transfection to enhance survival after transfection. After 24 hr of transfection, cells were washed once with sterile PBS and fixed with 4% formaldehyde solution (ref. 15710; Electron Microscopy Sciences, Hatfield, PA, USA) for 15 min at room temperature followed by three washing steps with PBS. When no staining was required, coverslips were directly mounted onto slides using Vectashield mounting medium without DAPI (Vector Laboratories, Burlingame, CA). For endosomal marker staining, fixed coverslips were permeabilized with 0.2% Triton X-100 (Sigma-Aldrich) for 10 minutes at room temperature and blocked with blocking solution [5% (w/v) BSA (ref. 160069, MP Biomedicals), 0.1% Triton X-100 in PBS] for one-hour at room temperature, then coverslips were incubated overnight at 4°C with antibodies diluted 1:200 in blocking solution (Endosomal Marker Antibody Sampler Kit #12666, Cell Signaling Technology). The following day, coverslips were washed three times with PBS and incubated with secondary antibodies for 1 hour at room temperature (1:1000 dilution, Alexa fluor 488 or 594 labelled highly cross-adsorbed goat anti-mouse or goat anti-rabbit secondary antibodies, Invitrogen, refs. A11029, A32731, A11032, A11037). Following additional three washing steps with PBS, coverslips were mounted onto slides using Vectashield. If DAPI staining was carried out, after the final washing steps with PBS, DAPI was added at 300nM concentration and incubated at room temperature for 5 minutes. Then, coverslips were washed additional two times with PBS and mounted to slides with Vectashield. Images were taken using a Zeiss LSM 710 confocal microscope with an 63x oil objective using excitation at 405 nm, 488 nm, and 561 nm and emission at 445-470, 500-550, and 570-610 nm for DAPI/CFP, GFP, and mCherry, respectively. Image analysis was done using Fiji ImageJ 1.53 c software.

### Single Molecule Pulldown

Coverslips were treated with ∼0.05% (v/v) dichlorodimethylsilane (DDS; Aldrich, >99.5%), then coated with neutravidin (0.2 mg/mL; Thermo Fisher) in T50 buffer for 5 min. Coverslips were then passivated with 0.2% Tween-20 (Fisher BioReagents) in T50 buffer for 10 min. Biotinylated 4-1BB bait variants (1 µM) in T50 buffer supplemented with BSA (0.1 mg/mL) and preincubated with TCEP (0.1%) were then applied for 10 min, followed by several washes with T50 + BSA. Coverslips were incubated with GFP-LANCL2 lysate (same as in nHU experiments) diluted in T50 + BSA for 10 min and washed repeatedly with T50 + BSA to minimize nonspecific binding. Single-molecule complexes were imaged using total internal reflection fluorescence (TIRF) microscopy (488 nm argon laser) with a 100× objective (Nikon). Movies (13 × 13 µmZ) of 500–1000 frames were acquired at 12.5 Hz and analyzed using Fiji (NIH). In total, 5 movies were captured for WT and C243S with ∼ 1000 and ∼700 molecules per movie, respectively, and 3 movies for C253S with ∼200 molecules per movie due to weak binding. Three movies were analyzed for photobleaching experiments with 30 particles per movie and the number of observed steps were counted (Supplemental Data 3).

### GSH/GSSG assay

The day before transfection, 3×10^4^ HEK293T cells/well were seeded on 96-well clear bottom white plates (Revvity ViewPlate-96 #6005181). Cells were co-transfected with 0.1 µg of total plasmid DNA per well using JetPRIME reagent (Polyplus, #101000046) according to the manufacturer’s recommendations, with equal amounts of each plasmid in co-transfections (0.033 µg/each plasmid). Each well was transfected with GFP-LANCL2, CFP-TRAF2 and a 4-1BB-mCherry variant except the mock controls that were only treated with the transfection reagent. The medium was changed to fresh medium after 5 hr of transfection to enhance survival. After 24 hr of transfection, the cellular GSH and GSSG relative levels or their ratio was determined using the GSH/GSSG-Glo™ Assay (Promega #V6611), which was carried out according to the manufacturer’s recommendations. Luminescence measurements for GSSG and total glutathione were carried out by a Molecular Devices SpectraMax iD5 multimode plate reader and final ratios were calculated from these relative luminescence units. In total, six biological replicates (six GSSG and six total glutathione values) were measured for each condition.

### NF-κB activity reporter assay using the 293T-kb-luc

Freshly thawed HEK293T cells were used for stable cell line generation. The optimal puromycin concentration was determined by a kill curve performed before stable selection. The cells were seeded in poly-D-lysine (PDL)-coated 24-well plates at 0.75 × 10^5^ cells per well. The following day, different amounts of puromycin (0-8 µg/mL) were added to the wells, and cells were incubated for 4 days. Viable and non-viable cells were quantified on day 4 using an EVE cell counter with trypan blue staining. Cell viability decreased in a dose-dependent manner, with complete cell death at 3 µg/mL. Finally, 2 µg/mL was chosen for selection. For stable cell line generation, a previously established Sleeping Beauty transposon system was used ^72,73^. Sleeping Beauty transposase containing pSB100 vector (Addgene #34879) was a kind gift of Dr. Zsuzsanna Izsvak (MDC Berlin Germany) and Dr. Tamas I. Orban (RCNS Budapest Hungary) ^74^. In the p10-CAG-MCS-IRES-GFP plasmid, the GFP cassette was replaced with a puromycin resistance cassette. A kb-luc construct previously used in transient assays (kind gift of the lab of Simona Saccani ^75^) containing three κB sites upstream of a chicken conalbumin promoter driving Firefly luc2 was cloned into this plasmid by replacing the CAG-MCS region downstream of the left inverted repeat/direct repeat (IR/DR-L) element. The resulting p10-kb-luc-IRES-PuroR plasmid contained both the luciferase reporter and a puromycin resistance cassette for selection.

For stable cell line generation, 1.5 × 10^6^ cells were seeded in 100 mm dishes pretreated with PDL. After 24 h, cells were transfected with the p10-kb-luc-IRES-PuroR construct together with the SB100 transposase plasmid at a 10:1 (p10:pSB100) ratio, using JetPrime reagent according to the manufacturer’s instructions. Non-transfected cells and cells transfected with p10 alone were used as controls. The following day, cells were allowed to recover, and puromycin selection (2 µg/mL) was initiated on day 2 post-transfection. Fresh selection medium was replaced every 2–3 days. On day 6, cells were transferred to CellBIND T75 flasks, and on day 8 to PDL-coated T175 flasks. A second wave of cell death was observed during this period, corresponding to elimination of non-stably integrated cells. Selection was maintained until day 13 at 2 µg/mL puromycin, after which the concentration was reduced to 0.5 µg/mL for maintenance. Stable cells were kept under this condition until subsequent experiments. Puromycin was withdrawn one week prior to transfection to enhance cell growth. Genomic integration of the transposon was validated by quantitative PCR (qPCR) on genomic DNA isolated using Qiagen FlexiGene DNA Kit (Ref. #51206). As a reference locus, we used the single-copy gene RPPH1, located on chromosome 14q11.2. Although HEK293T cells are aneuploid and near-triploid, chromosome 14 is generally considered to be present in two copies in this cell line and was therefore used as a diploid reference ^76^. The IR/DR-L sequence of the transposon was amplified to quantify transposon integration events (as done before ^73^). qPCR was performed using a QuantStudio 3 Real-Time PCR System (Thermo Fisher Scientific) and SYBR Green PowerTrack Master Mix (Ref. #A46109, Thermo Fisher Scientific). Comparison of the IR/DR-L and RPPH1 amplification signals revealed an approximately 10-fold higher abundance of the transposon sequence relative to the diploid reference locus. Assuming two copies of RPPH1 per cell, this corresponds to an estimated average of approximately 20 integrated transposon copies per cell.

For transfection, stable cells were plated on a 96-well clear bottom white plates (Revvity ViewPlate-96 #6005181) at 1.5×10^4^ cells/well concentration. The next day, in total 0.1 µg DNA (0.033 µg/each plasmid) was transfected to the cells using JetPRIME reagent (Polyplus) according to the manufacturer’s recommendations. We used the following constructs to transfect the cells: pCEP4-mCherry or pCEP4-4-1BB-wt-mCherry; pEGFP-C1 or pEGFP-LANCL2; pECFP-C1 or pECFP-TRAF2. In case of mock, only transfection reagent was used during transfection. The medium was replaced after 5 hours to enhance cellular survival.

24 hours after transfection, cellular luminescence was measured using Promega Steady-Glo Luciferase Assay System (#E2510) according to the manufacturer’s instructions. Briefly, we removed the cells from the incubator 15 minutes before the measurement to equilibrate at RT. We added 100 µl of Steady-Glo reagent to each wells containing 100 µl medium and briefly mixed the plate. After 5 minutes of waiting for cellular lysis, luminescence (relative light unit) was measured in a Molecular Devices SpectraMax iD5 multimode plate reader. In total, 6 biological replicates were measured for each condition.

### RNA expression analysis of NF-κB target genes

HEK293T cells were seeded on standard 24-well plates (Costar #3526) at 1.5×10^5^ cells/well concentration. The next day, cells were transfected with the different constructs (pCEP4-mCherry or pCEP4-4-1BB-wt-mCherry; pEGFP-C1 or pEGFP-LANCL2; pECFP-C1 or pECFP-TRAF2) using JetPRIME reagent according to the manufacturer’s recommendations. We used in total 0.45 µg DNA (0.15 µg/each plasmid) and 1 µl JetPRIME reagent/well. After 5 hours, the medium was replaced to enhance cell survival. RNA was isolated after 28-30 hours of transfection using Macherey-Nagel Nucleo-Spin RNA Kit (#740955). Concentration of RNA was determined using a Thermo Scientific NanoDrop One instrument. Reverse transcription was performed using the same amounts of RNA (generally 1.5 ug/each condition) and SuperScript IV RT kit (Thermo Fisher Scientific #18090200) according to the manufacturer’s instructions with random hexamers (Thermo Fisher Scientific #N8080127) and RNasin (Promega #N2511). Quantitative real-time PCR was performed in a QuantStudio 3 instrument (Applied Biosystems) using PowerTrack SYBR Green Master Mix (Thermo Fisher Scientific #A46109) with the following primers: NFKBIA_for: 5’-TCCACTCCATCCTGAAGGCTAC-3’, NFKBIA_rev: 5’-CAAGGACACCAAAAGCTCCACG-3’, IL8_for: 5’-GAGAGTGATTGAGAGTGGACCAC-3’, IL8_rev: 5’-CACAACCCTCTGCACCCAGTTT-3’, TNFAIP3_for: 5’-CTCAACTGGTGTCGAGAAGTCC-3’, TNFAIP3_rev: 5’-TTCCTTGAGCGTGCTGAACAGC-3’, RPLP0_for: 5’-TGGTCATCCAGCAGGTGTTCGA-3’, RPLP0_rev: 5’-ACAGACACTGGCAACATTGCGG-3’. Standard fast cycling experiment method was used: Initial activation 95°C for 20 sec, Cycling 40x: Denaturation 95°C for 1 sec and annealing/extension/fluorescence acquisition 60°C for 20 sec. Relative expression levels (fold changes) were determined using standard 2^-ΔΔCT^ method by normalizing to reference gene RPLP0 values (ΔCT) and non-transfected controls (ΔΔCT). In total, three biological replicates were performed, each including three technical replicates.

### Statistics

Two-tailed unpaired T-test, assuming equal variance was used to evaluate mass spectrometry data and GSH/GSSG levels. Two-tailed unpaired T-test was performed for statistics and Bonferroni correction was applied for NFκB activity experiments because of multiple testing was performed on various conditions.

## Data availability

Raw mass spectrometry data measured on the Exploris and timsTOF systems are available via ProteomeXchange with identifiers PXD085745 and PXD086181, respectively. Affinity interactomic data is also released on the ProfAff database (profaff.fr).

## Supplementary Figures

**Supplemental Figure 1.**
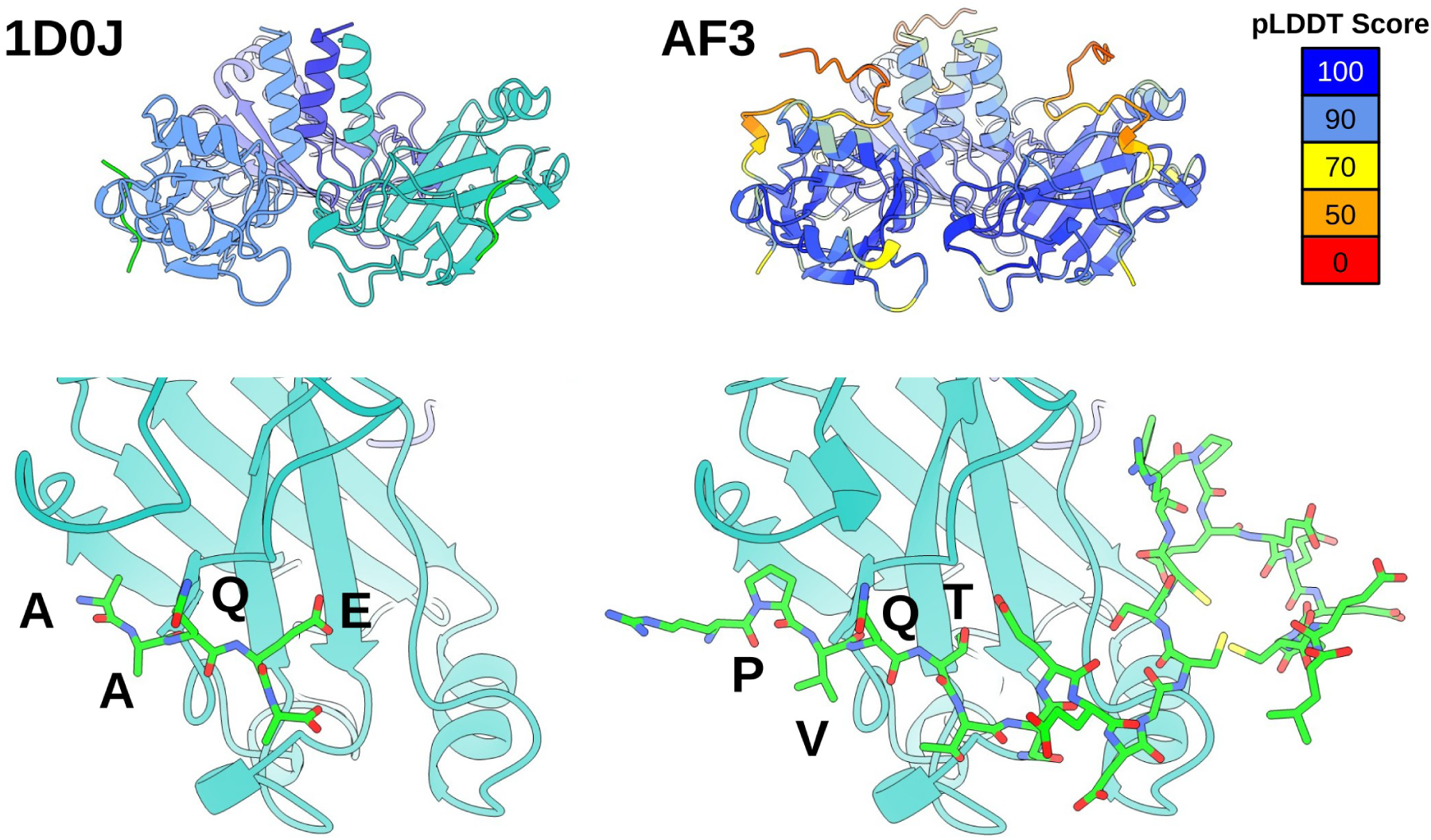
(left) Crystal structure of the complex between (mouse) 4-1BB and TRAF2. (right) AlphaFold3 prediction with default settings using the core trimeric TRAF2 domain and the entire cytoplasmic tail of 4-1BB. Upper panel is colored by local confidence scores. Only the core PVQT was predicted confidentally. Peptide is shown in green. The overview image for the predicted structure is colored based on pLDDT scores.

**Supplemental Figure 2.**
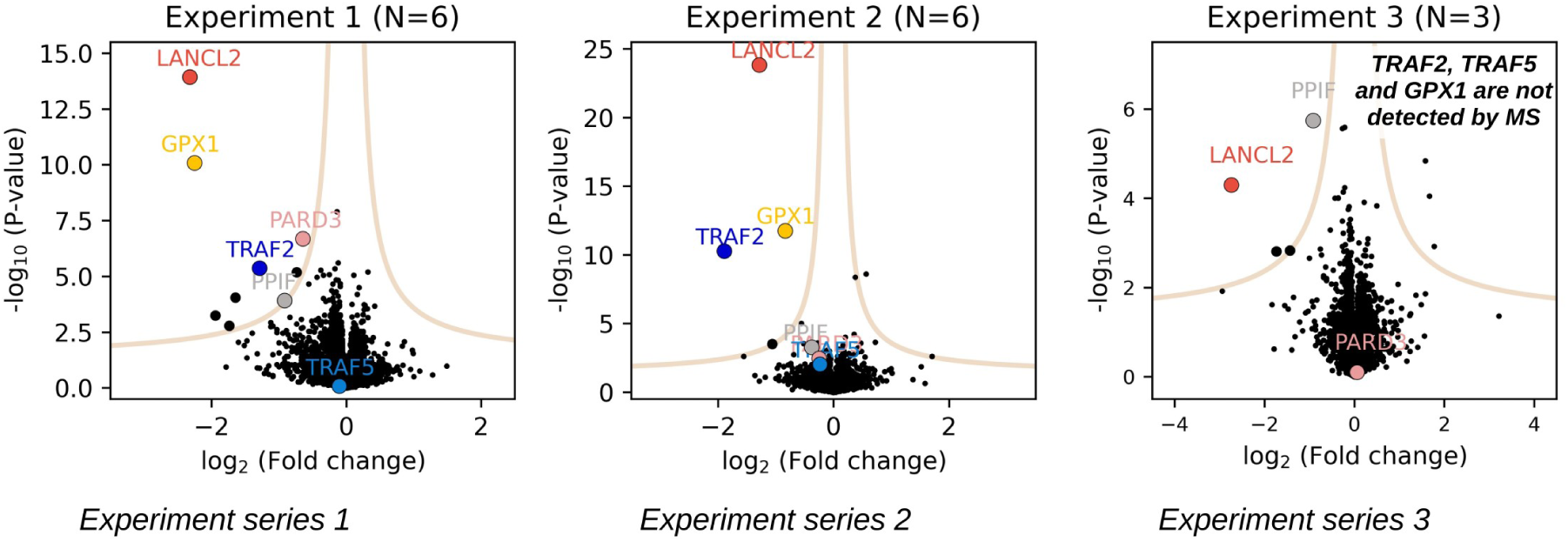
Individual nHU-MS experiments on the 4-1BB tail peptide using Jurkat cell extracts. Note that TRAF2, TRAF5 and GPX1 are not detected in the experiment series 3.

**Supplemental Figure 3.**
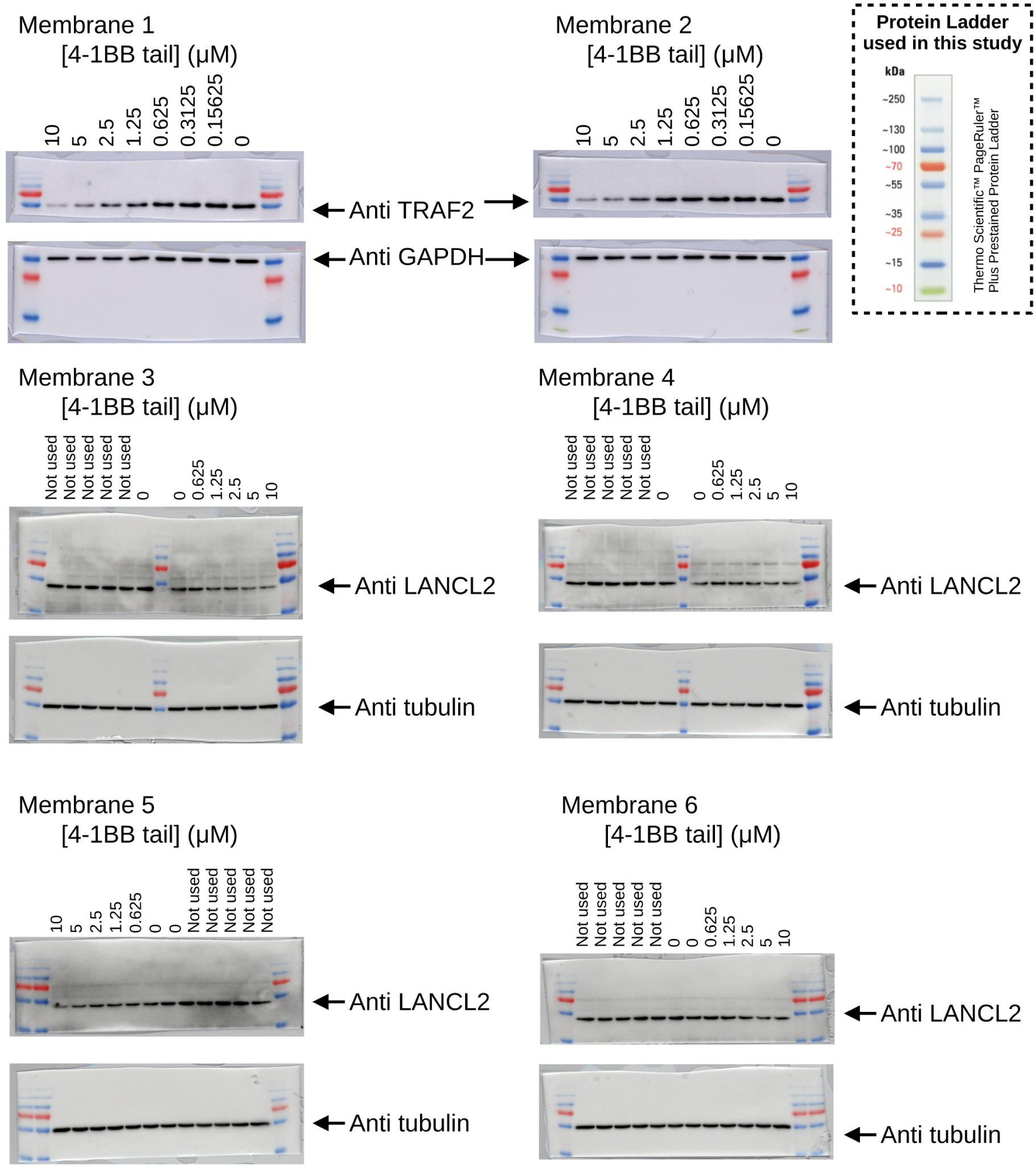
nHU-WB experiments showing uncropped blots with overlay colorimetric signal.

**Supplemental Figure 4.**
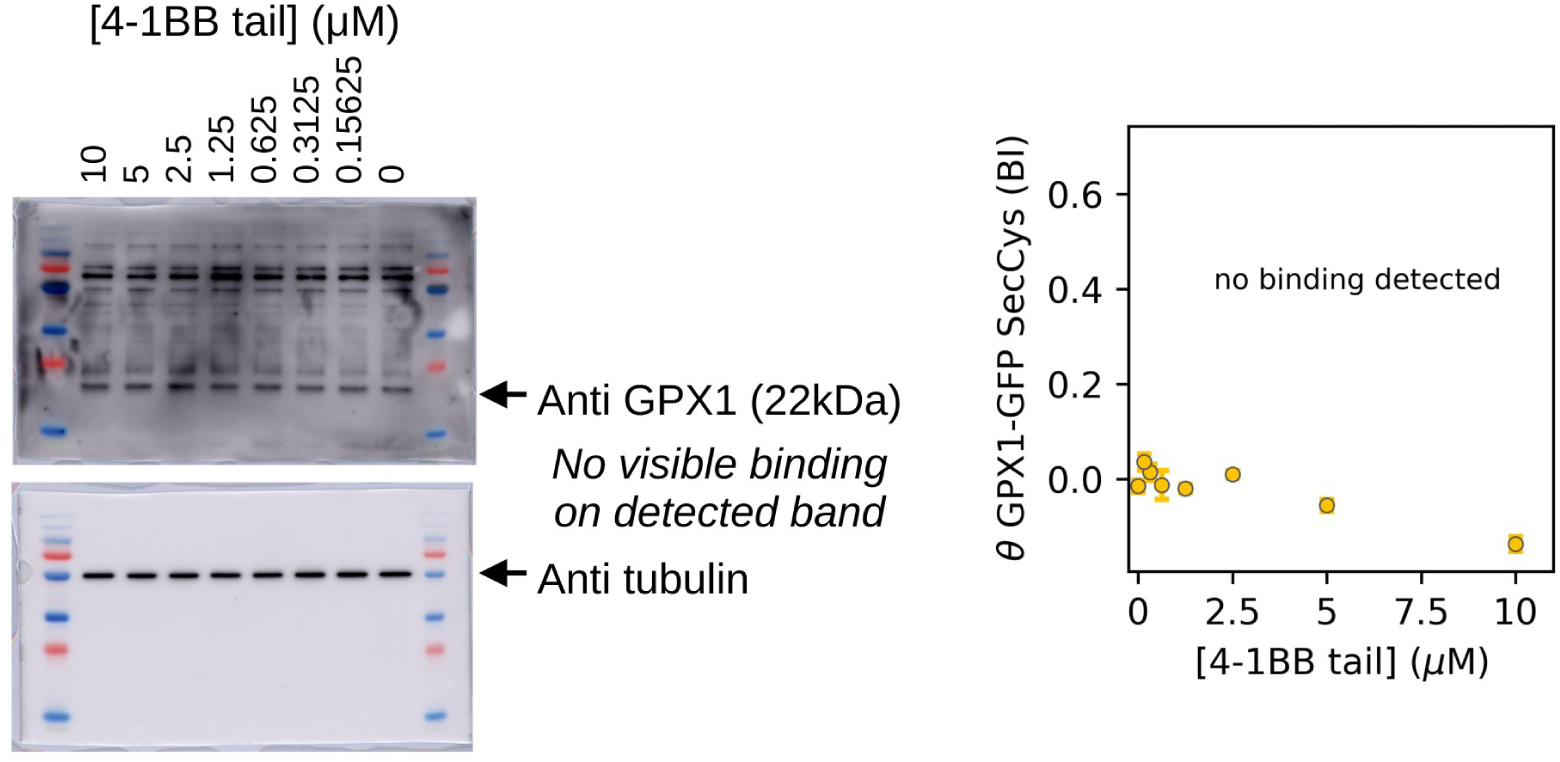
nHU-WB and nHU-F experiments on GPX1. (left) nHU-WB experiment on endogenous GPX1 showing uncropped blots with overlay colorimetric signal. No visible band was identified at the correct molecular weight that showed binding to 4-1BB. (right) nHU-F measurement with GFP-tagged GPX1, with a Sec mutated to Cys does not show binding to 4-1BB.

**Supplemental Figure 5.**
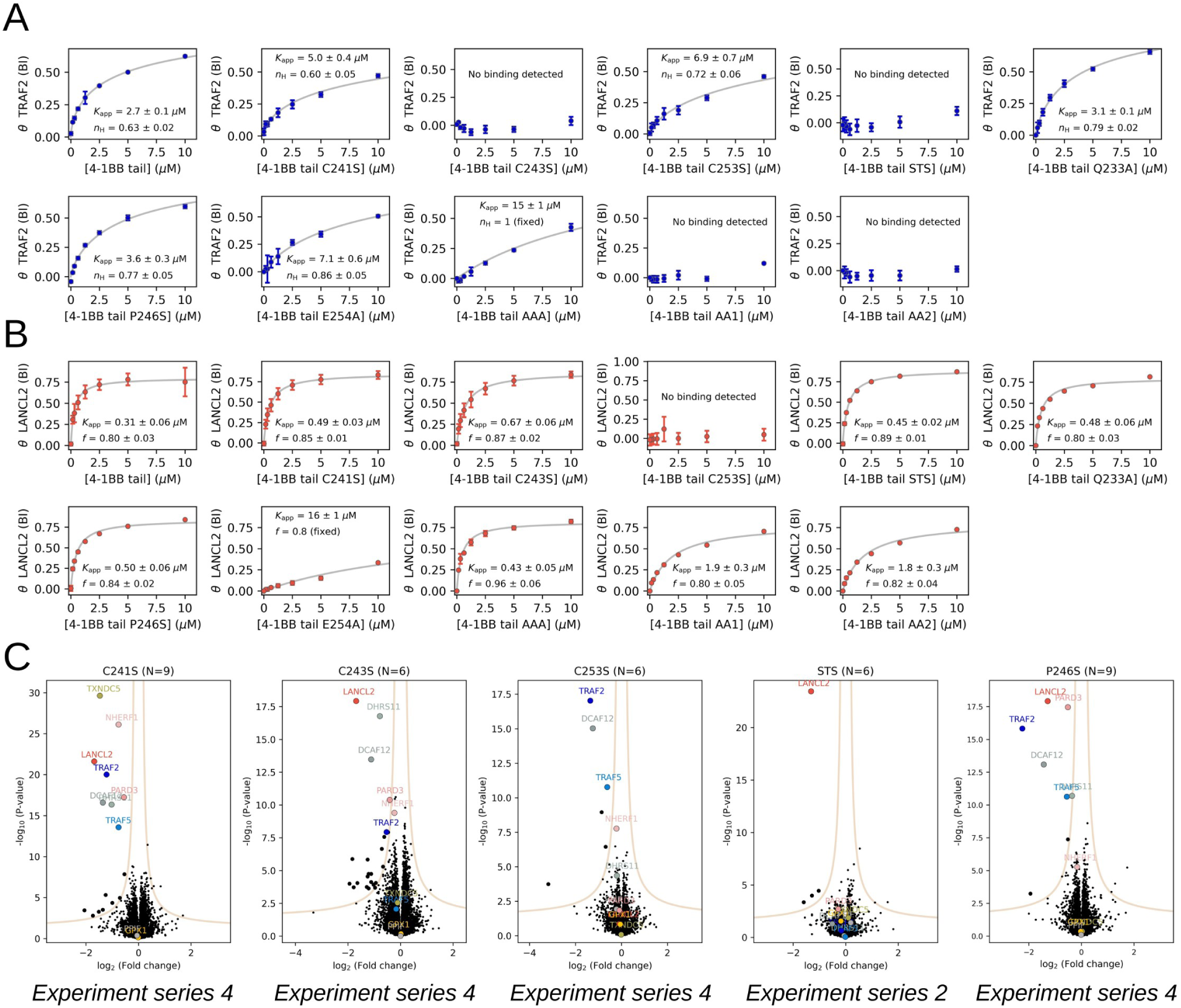
Interactomes of 4-1BB variants. (A) nHU experiments with GFP-fused TRAF2, measured by fluorescence intensity. Each 8-point binding experiment was performed with a different 4-1BB variant and fluorescence intensity was measured in technical triplicates. (B) nHU experiments with GFP-fused LANCL2, measured by fluorescence intensity. Each 8-point binding experiment was performed with a different 4-1BB variant and fluorescence intensity was measured in technical triplicates. (C) Proteome-wide nHU experiments on selected 4-1BB mutants show changes in the interactome, in agreement with the focused experiments.

**Supplemental Figure 6.**
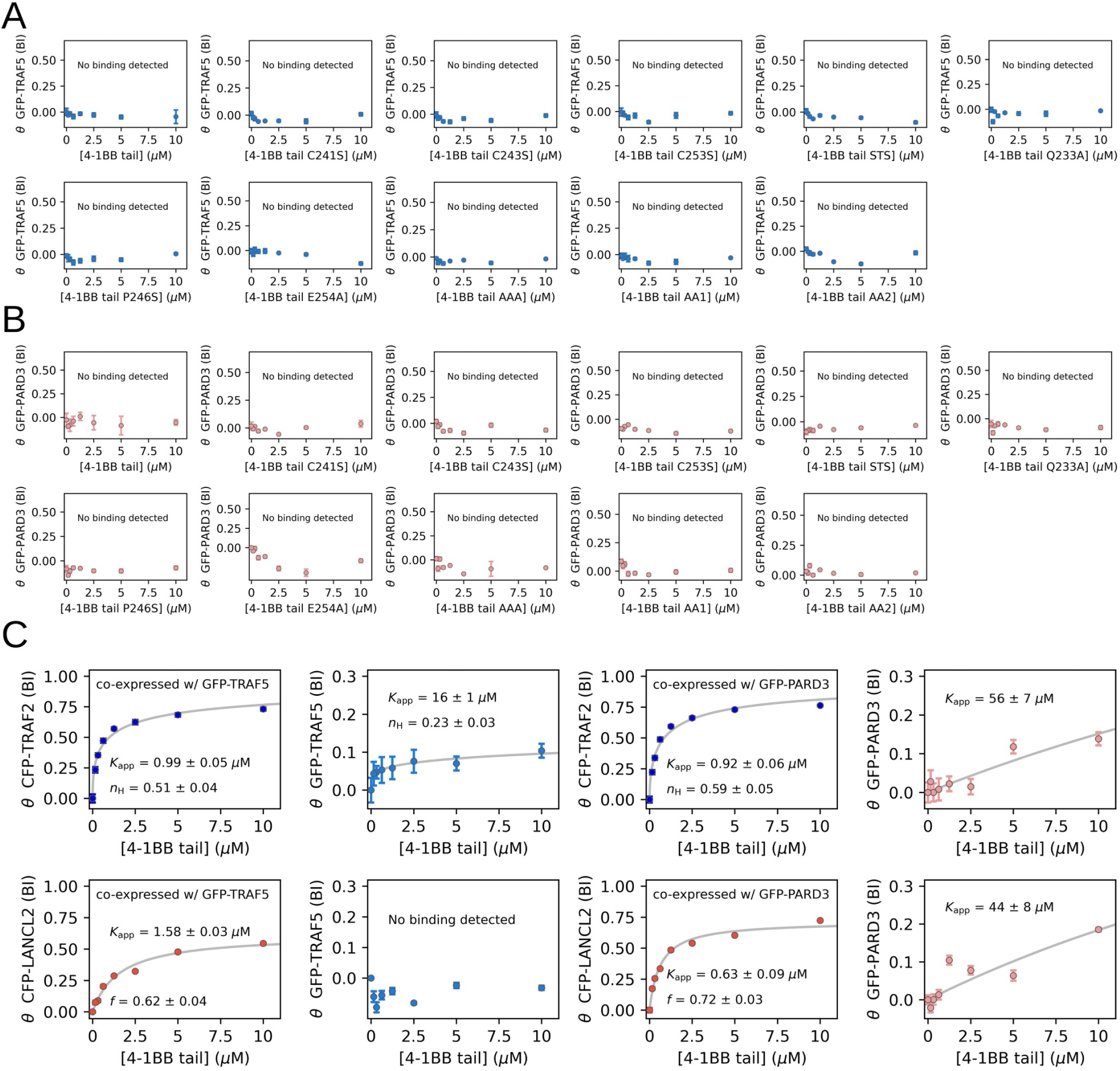
Interaction between 4-1BB and TRAF5 or PARD3. (A) nHU experiments with GFP-fused TRAF5, measured by fluorescence intensity. Each 8-point binding experiment was performed with a different 4-1BB variant and fluorescence intensity was measured in technical triplicates. None of the 4-1BB variants display binding to TRAF5. (B) nHU experiments with GFP-fused PARD3, measured by fluorescence intensity. Each 8-point binding experiment was performed with a different 4-1BB variant and fluorescence intensity was measured in technical triplicates. None of the 4-1BB variants display binding to PARD3. (C) nHU experiments with GFP-fused TRAF5 or PARD3, coexpressed with CFP-fused TRAF2 or LANCL2, measured by fluorescence intensity. Each 8-point binding experiment was performed with a different 4-1BB variant and fluorescence intensity was measured in technical triplicates. TRAF5 shows binding in the presence of TRAF2. PARD3 shows marginal binding in the presence of either TRAF2 or LANCL2. Experimental pairs with co-expressed proteins are shown next to each other.

**Supplemental Figure 7.**
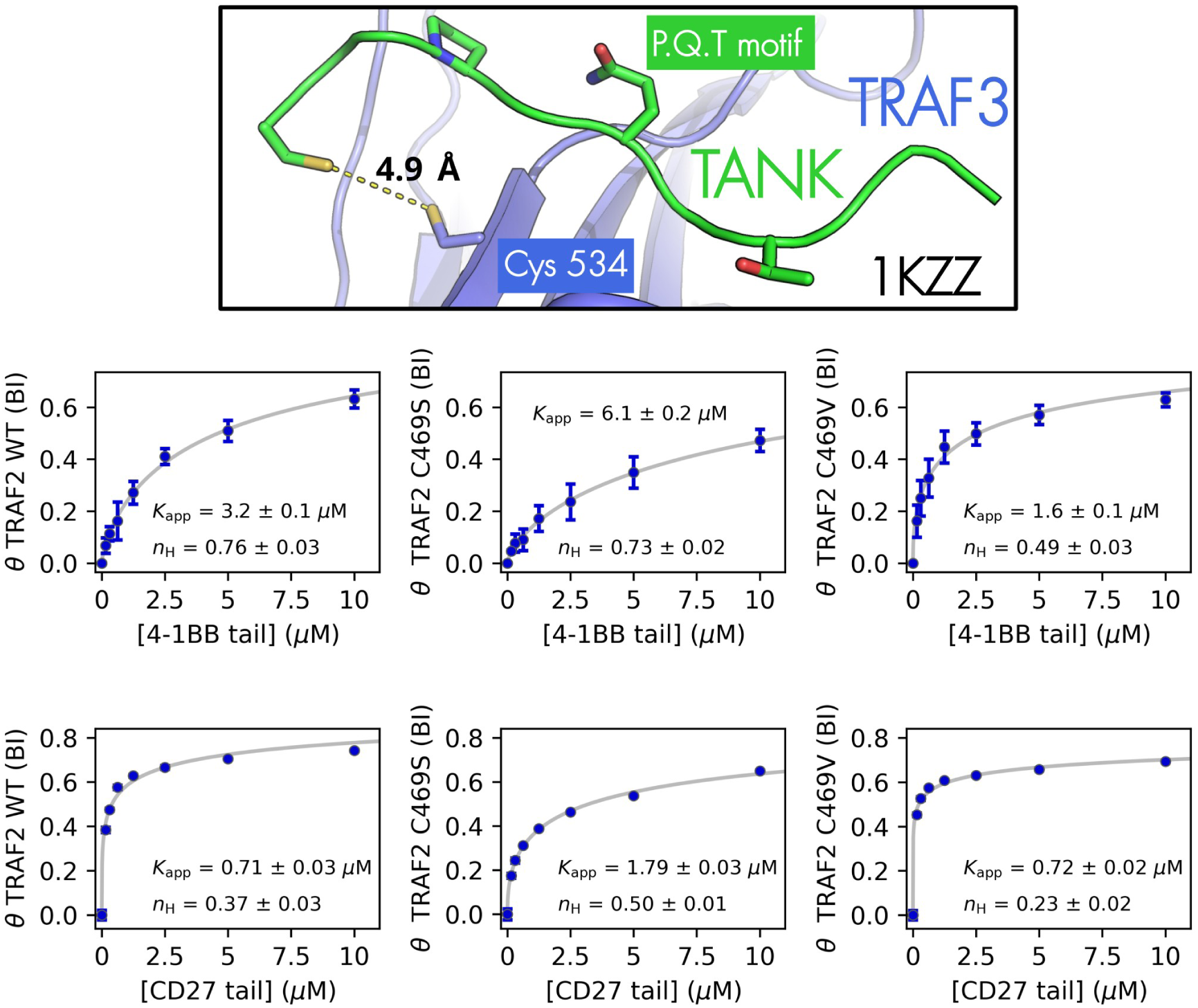
Cys-dependent exosite on TRAF proteins. (up) Crystal structure of the complex of TRAF3 and a peptide motif from TANK shows the close proximity of a Cys from upstream the binding site and Cys 534 from TRAF3. (down) nHU experiments with GFP-fused TRAF2 mutants against 4-1BB or CD27, measured by fluorescence intensity. Each 8-point binding experiment was performed with a different TRAF2 variant and fluorescence intensity was measured in technical triplicates.

**Supplemental Figure 8.**
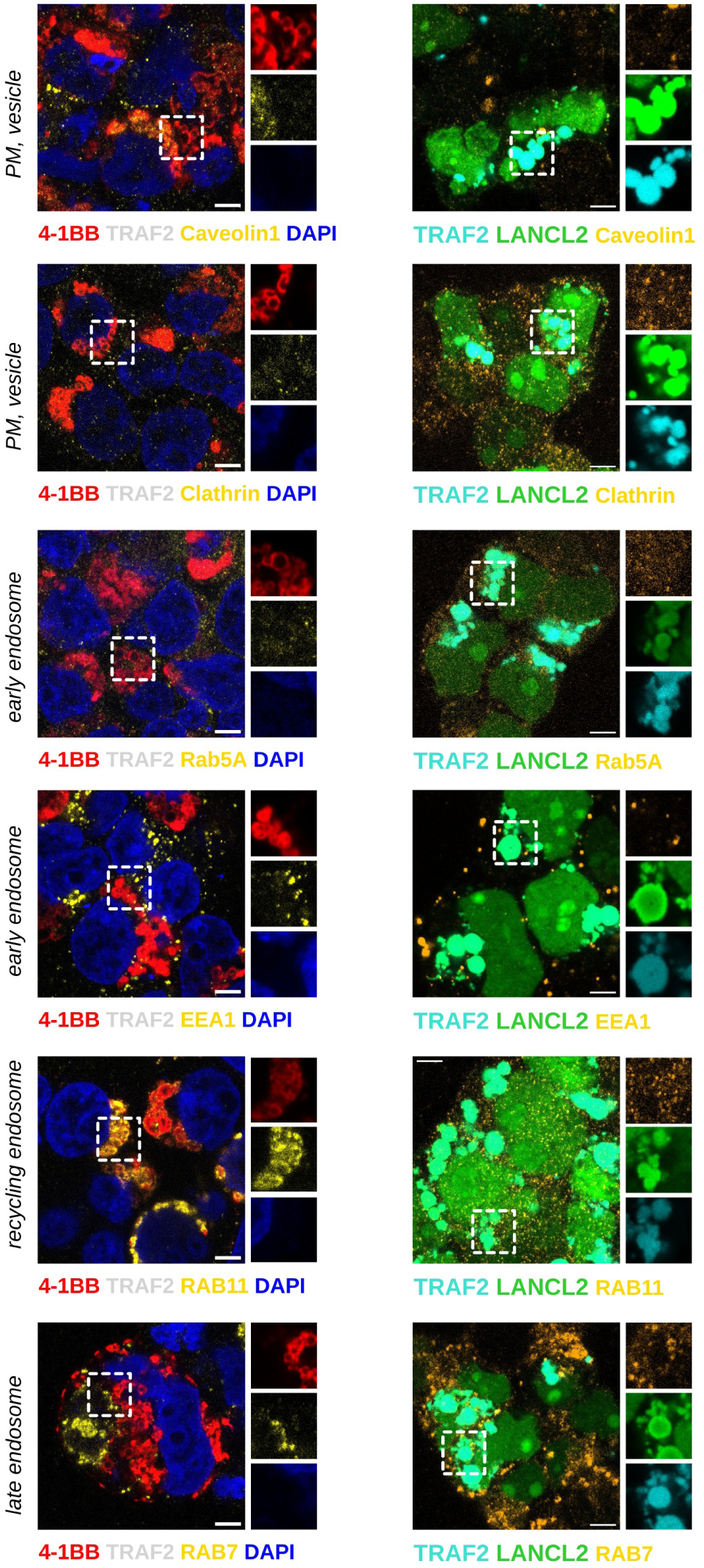
Co-localization of 4-1BB, or TRAF2, and LANCL2 with endosomal markers. Confocal images were taken from fixed HEK293T cells transfected with fluorescently tagged proteins. (left) Fluorescence images of 4-1BB and endosomal markers (shown in yellow, stained with Alexa 488-labelled secondary antibody) on HEK293T cells transfected with 4-1BB-mCherry and untagged TRAF2. DAPI staining is shown in blue. Only RAB11 shows clear co-localization with 4-1BB. (right) Fluorescence images of CFP-TRAF2 and GFP-LANCL2 and endosomal markers (shown in yellow, stained with Alexa 594-labelled secondary antibody) on transfected HEK293T cells For each panel, an overlay signal is shown on the left and the insets on the right show separate channels. Scale bar is equivalent to 5 μm. Representative images are shown based on hundreds of observations.

**Supplemental Figure 9.**
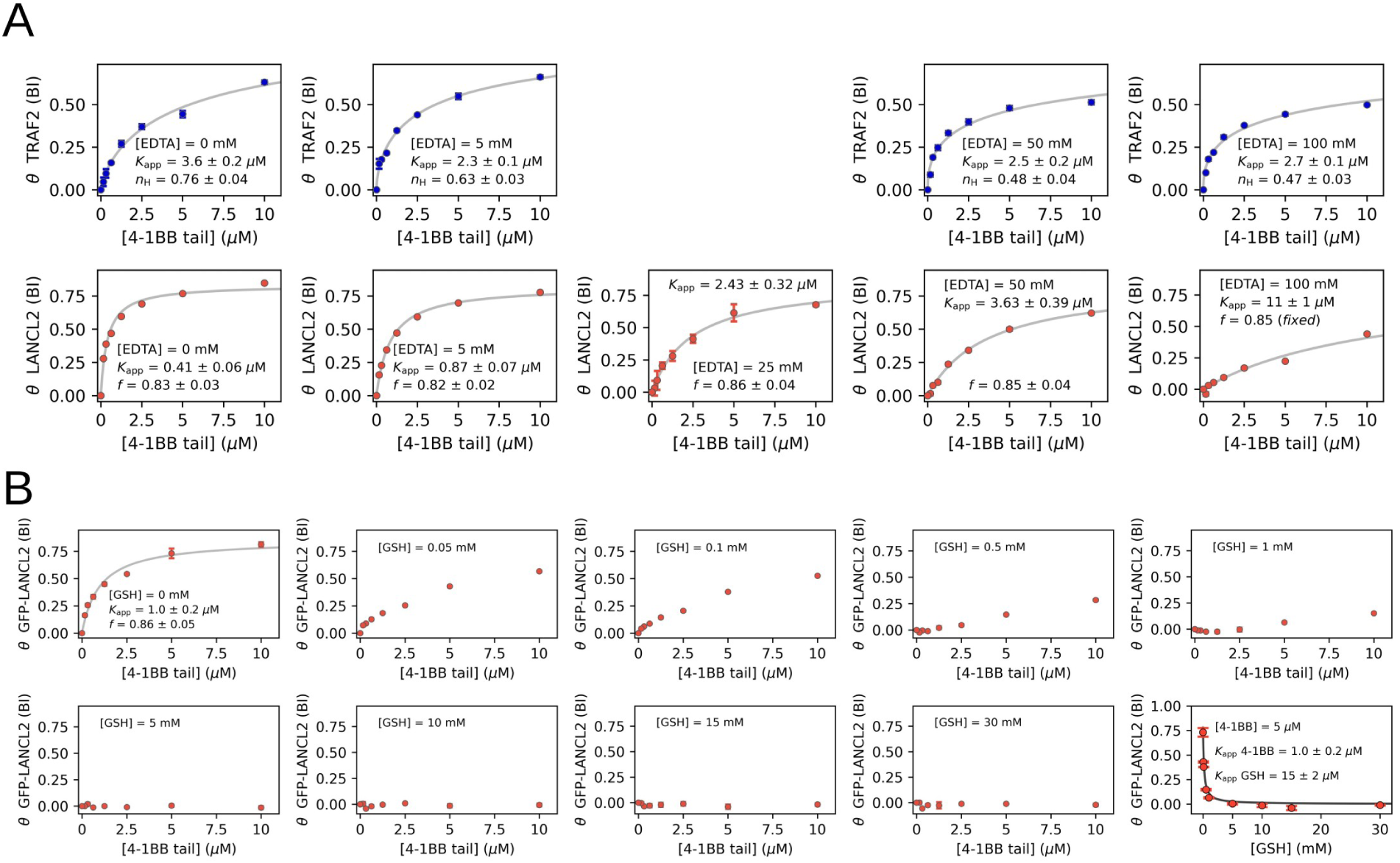
LANCL2 binding is disrupted by EDTA and competes with glutathione. (A) nHU experiments with GFP-fused TRAF2 or LANCL2 against 4-1BB in the presence of increasing amount of EDTA, measured by fluorescence intensity. Each 8-point binding experiment was performed with a different EDTA concentration and fluorescence intensity was measured in technical triplicates. (B) nHU experiments with GFP-fused LANCL2 against 4-1BB in the presence of increasing amount of reduced glutathione, measured by fluorescence intensity. Each 8-point binding experiment was performed with a different GSH concentration and fluorescence intensity was measured in technical triplicates.

**Supplemental Figure 10.**
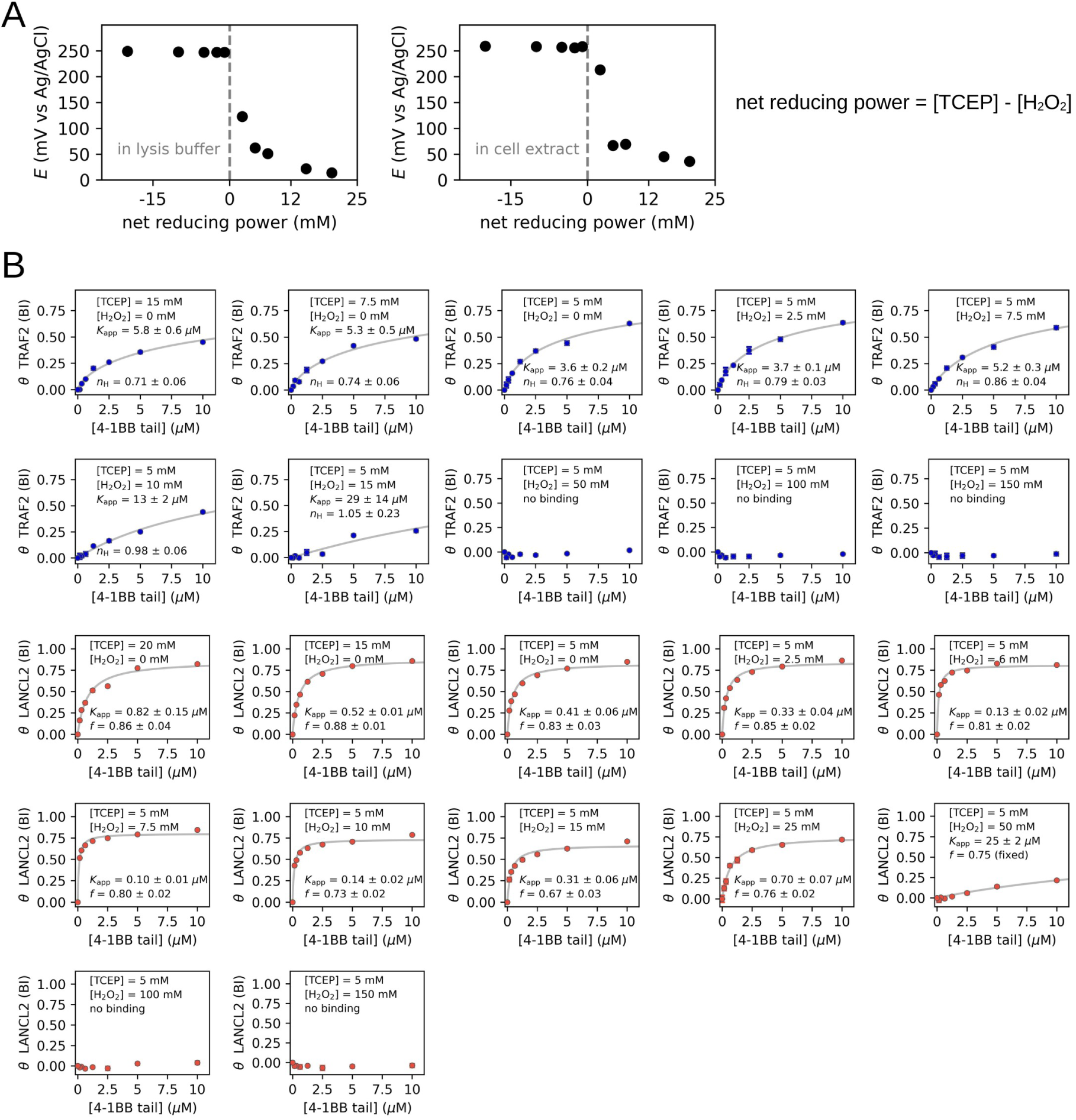
TRAF2 and LANCL2 interactions with 4-1BB shows redox-dependence (A) Potentiometric measurements of redox potential in mixtures of H2O2 and TCEP in lysis buffer or in total cell lysate (with 0.1 mg/ml total protein content). (B) nHU experiments with GFP-fused TRAF2 or LANCL2 against 4-1BB in the presence of various amount of H_2_O_2_ and TCEP, measured by fluorescence intensity. Each 8-point binding experiment was performed with a different H_2_O_2_ and TCEP concentrations and fluorescence intensity was measured in technical triplicates.

**Supplemental Figure 11.**
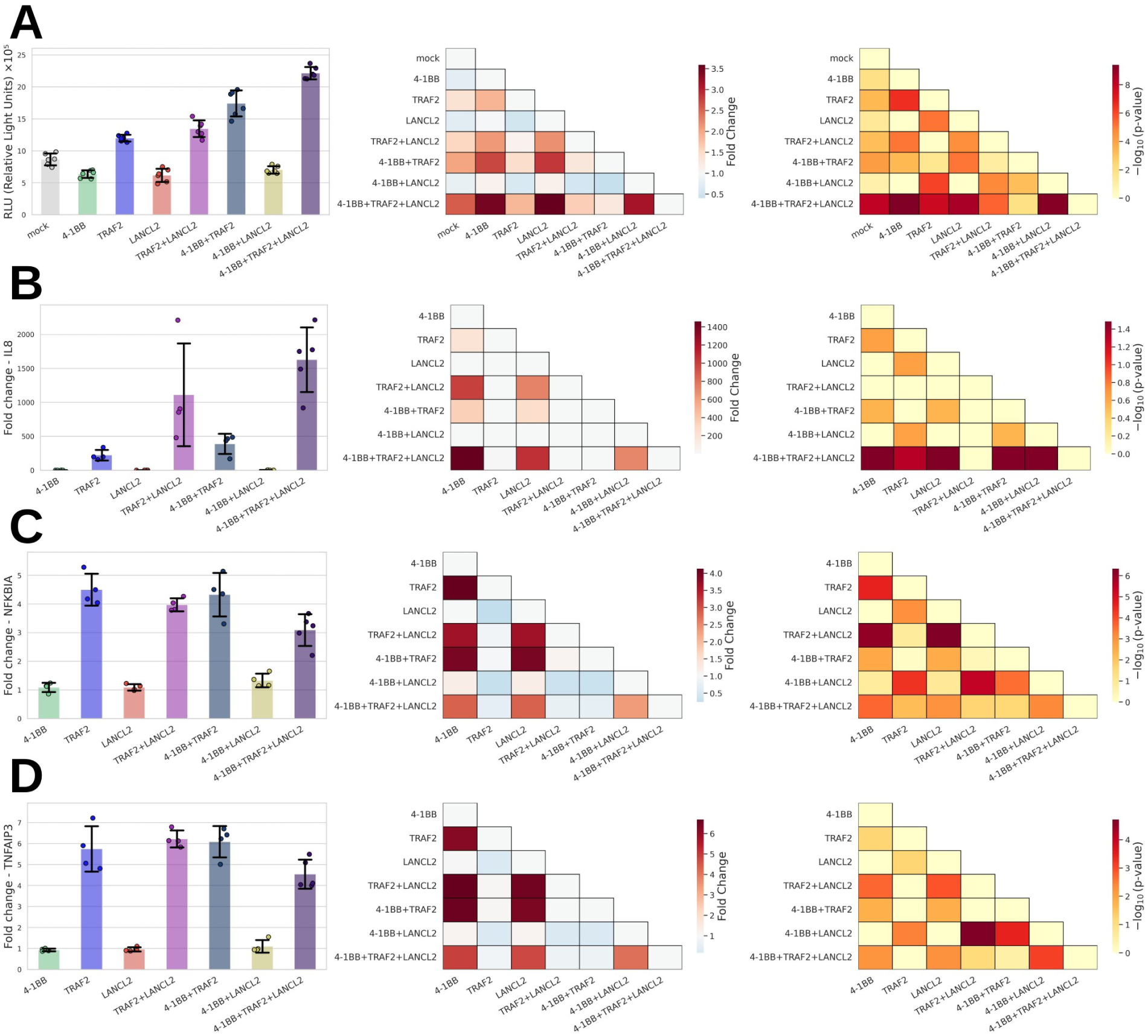
LANCL2 affects NFκB activity. (A) A genetic kb-luc-based assay was developed to monitor NFκB activity in 293T cells transfected with 4-1BB, TRAF2, LANCL2, or their combinations. (B-D) Transcription levels of known NFκB targets were measured in 293T cells transfected with 4-1BB, TRAF2, LANCL2, or their combinations. The averages of technical triplicates are shown from at least 3 biological replicates. Two-tailed unpaired T-test was performed for statistics and Bonferroni correction was applied because of multiple testing. The data of first panels of A, B, C are identical to those found in Figure 6 and kept here only for clarity.

## Supplemental data

**Supplemental Data 1**, Summary of nHU-MS experiments and proteomic/stoichiometric calculations.

**Supplemental Data 2**, Summary of titration nHU experiments.

**Supplemental Data 3**, Summary of SiM-Pull experiments.

## Notes

### Competing Interest Statement

The authors have declared no competing interest.

